# Defining ancestry, heritability and plasticity of cellular phenotypes in somatic evolution

**DOI:** 10.1101/2022.12.28.522128

**Authors:** Joshua S. Schiffman, Andrew R. D’Avino, Tamara Prieto, Yakun Pang, Yilin Fan, Srinivas Rajagopalan, Catherine Potenski, Toshiro Hara, Mario L. Suvà, Charles Gawad, Dan A. Landau

**Affiliations:** New York Genome Center, New York, NY, USA; Weill Cornell Medicine, New York, NY, USA; Tri-Institutional MD-PhD Program, Weill Cornell Medicine, Rockefeller University, Memorial Sloan Kettering Cancer Center, New York, NY, USA; Stanford University, Stanford, CA, USA; St. Jude Children’s Research Hospital, Memphis, TN, USA; Department of Pathology and Center for Cancer Research, Massachusetts General Hospital and Harvard Medical School, Boston, MA, USA; Broad Institute of Harvard and MIT, Cambridge, MA, USA; Chan Zuckerberg Biohub, San Francisco, CA, USA

## Abstract

The broad application of single-cell RNA sequencing has revealed transcriptional cell state heterogeneity across diverse healthy and malignant somatic tissues. Recent advances in lineage tracing technologies have further enabled the simultaneous capture of cell transcriptional state along with cellular ancestry thus enabling the study of somatic evolution at an unprecedented resolution; however, new analytical approaches are needed to fully harness these data. Here we introduce PATH (Phylogenetic Analysis of Transcriptional Heritability), an analytical framework, which draws upon classic approaches in species evolution, to quantify heritability and plasticity of somatic phenotypes, including transcriptional states. The PATH framework further allows for the inference of cell state transition dynamics by linking a model of cellular evolutionary dynamics with our measure of heritability versus plasticity. We evaluate the robustness of this approach by testing a range of biological and technical features in simulations of somatic evolution. We then apply PATH to characterize previously published and newly generated single-cell phylogenies, reconstructed from either native or artificial lineage markers, with matching cellular state profiling. PATH recovered developmental relationships in mouse embryogenesis, and revealed how anatomic proximity influences neural relatedness in the developing zebrafish brain. In cancer, PATH dissected the heritability of the epithelial-to-mesenchymal transition in a mouse model of pancreatic cancer, and the heritability versus plasticity of transcriptionally-defined cell states in human glioblastoma. Finally, PATH revealed phenotypic heritability patterns in a phylogeny reconstructed from single-cell whole genome sequencing of a B-cell acute lymphoblastic leukemia patient sample. Altogether, by bringing together perspectives from evolutionary biology and emerging single-cell technologies, PATH formally connects the analysis of cell state diversity and somatic evolution, providing quantification of critical aspects of these processes and replacing *qualitative* conceptions of “plasticity” with *quantitative* measures of cell state transitions and heritability.

## Introduction

The application of single-cell RNA sequencing (scRNAseq) across biology has revealed vast phenotypic diversity within healthy [Hammond et al., 2019, Papalexi and Satija, 2018, Plasschaert et al., 2018] and diseased [Neftel et al., 2019, Wu et al., 2021] tissues. As genetic variation is limited within the soma, much of the heritable diversity of somatic phenotypes is attributed to non-genetic sources, such as epigenetic modifications. Indeed, the stable propagation of somatic phenotypes (*e.g*., cell type [Zeng, 2022]) through mitotic divisions, sometimes called *epigenetic memory* [Fennell et al., 2022, Halley-Stott and Gurdon, 2013, Larsen et al., 2021, Shaffer et al., 2020], often relies on the heritable transmission of epigenetic marks, such as DNA methylation, histone modification, or the propagation of key transcription factors [Adam and Fuchs, 2016, Whyte et al., 2013]. Somatic cells, however, may also accumulate genetic variation over time [Li et al., 2020, Martincorena et al., 2015, 2018], for example enabling more proliferative phenotypes that can lead to cancer [Hanahan, 2022, Vogelstein et al., 2013]. In addition to cell-intrinsic sources of heritable phenotypic diversity, cell-extrinsic sources, such as the microenvironment [Gola and Fuchs, 2021, Hara et al., 2021] or morphogen gradients [Houchmandzadeh et al., 2002], may contribute to heritable cellular phenotypic diversity, as progeny often share the same microenvironment as parent cells. Crucially, not all cellular phenotypic variation is stable, and cells can also plastically toggle between phenotypes in somatic evolution. For instance, healthy skin cells can dedifferentiate to repair injuries [Donati et al., 2017, Gola and Fuchs, 2021] and cancer cells have been shown to toggle between proliferative and invasive phenotypes [Karras et al., 2022, Oren et al., 2021], or to morph and evade treatment [Chan et al., 2022].

To approach these key aspects, it can be useful to consider cellular phenotypic diversity from an evolutionary perspective. Somatic cells descend from a common ancestor, and following successive divisions, accumulate heritable variation in the form of genetic, epigenetic or cell-extrinsic changes. Throughout this process of *somatic evolution*, the heritable variation within a population can be sculpted by selection, which has important implications for organismal health. Outcomes of somatic evolution, for instance, include the initiation, relapse, and treatment resistance of cancers [Fennell et al., 2022, Jan et al., 2012, Shaffer et al., 2017]. However, it is not yet clear to what degree epigenetic [Mazor et al., 2016] or genetic [Househam et al., 2022, Turajlic et al., 2019] variation contributes to the evolution and persistence of malignant phenotypes [Nam et al., 2021]. To confront the challenge of studying somatic evolution, we require an integrative model of somatic evolution that considers cellular phenotypic diversity and ancestry [Nam et al., 2021], informed by technologies that deliver phenotypically annotated single-cell phylogenetic trees [Biddy et al., 2018]. By tracing cellular ancestries, we can begin to elucidate the shared developmental origins of cell states and map differentiation trajectories [Chan et al., 2019, Raj et al., 2018]. Furthermore, this framework can enable us to dissect the heritability versus plasticity of somatic cellular phenotypes, to define how evolution shapes somatic cellular populations.

Recently, an array of techniques for lineage tracing has been advanced that can provide ancestry information at a singlecell level [Baron and van Oudenaarden, 2019, Sankaran et al., 2022]. In model organisms, cellular lineages or phylogenies can be reconstructed from *artificial* lineage markers [Pei et al., 2020, Raj et al., 2018, Rodriguez-Fraticelli et al., 2020, Spanjaard et al., 2018] that can be experimentally inserted and edited. In contrast, retracing lineage histories in human samples leverages *native* lineage markers, such as patterns of genetic (copy number [Salehi et al., 2022, Wang et al., 2021] or single nucleotide [Lodato et al., 2015, Ludwig et al., 2019]) or epigenetic (stochastic methylation [Gaiti et al., 2019]) variation. Both artificial and native lineage tracing approaches can be combined with other single-cell modalities, like scRNAseq, to deliver phylogenetic trees with phenotypically annotated leaves (terminal nodes).

Such phenotypically annotated cellular lineages emerge as a formidable tool to study critical questions in biology, such as mapping the ontogenetic relations between cells in development [Bandler et al., 2021], and clinically important features of cancer evolution, such as the stability of differentiation hierarchies [Chaligne et al., 2021], and metastatic dynamics [Quinn et al., 2021]. These experimental advances need to be complemented by a broadly applicable analytical framework, grounded in evolutionary biology, that could be applied to examine how cellular state (as for example profiled by scRNAseq) depends on ancestry (delivered by lineage tracing). Such a framework would enable us to distinguish between mitotically stable and ephemeral phenotypic states, and to make inferences about unobserved evolutionary dynamics. Tools for the analysis of multimodal single-cell lineages, such as *Hotspot* [Detomaso and Yosef, 2021] and *The Lorax* [Minkina et al., 2022], and others [Chaligne et al., 2021, Fang et al., 2022, Jones et al., 2022, Wang et al., 2022, Yang et al., 2022], are being developed to measure heritability. Nonetheless, additional conceptual and analytic advances are needed to fully harness these datasets for the study of somatic evolution. These advances will allow us to account for technical and biological variables affecting heritability measurements, and enable the integration of heritability assessments with phenotypic transition probability measurements, within a comprehensive and easy-to-implement analytical framework.

To address this challenge, we introduce **PATH** (**P**hylogenetic **A**nalysis of **T**ranscriptional **H**eritability), an analytical framework that draws upon classic approaches in species evolution, to quantify heritability and plasticity of somatic cellular phenotypes, such as transcriptional cell states. PATH measures *phylogenetic correlations*, which quantify the degree by which cellular phenotypes, broadly defined (*e.g*., transcriptional program, cell state or location), depend on ancestry, as provided by single-cell phylogenies, and *thus defines a measure of somatic heritability versus plasticity*. PATH builds upon auto-correlative [Cheverud and Dow, 1985, Gittleman and Kot, 1990] methods classically used to measure *phylogenetic signal* [Blomberg and Garland, 2002], the phylogenetic clustering of species phenotypes. Furthermore, PATH generalizes this approach to measure phylogenetic correlations *between* phenotypes (and from across modalities), providing a measure of how distinct phenotypes co-cluster on phylogenies, and thus defining a pairwise measure of phylogenetic signal. Additionally, for categorical phenotypes, such as cell type, PATH can transform phylogenetic correlations, our measurement of heritability versus plasticity, into inferences of transition rates between cell types or states. Importantly, this transformation provides a concrete interpretation of what phylogenetic signal measures, as the *pattern* of phylogenetic signal is directly linked with the *process* of cell type or state toggling. Further, PATH represents a comprehensive, versatile quantitative framework that can handle sparsely sampled and lowly resolved phylogenies, reconstructed under a range of biological and technical variables.

We first demonstrate PATH’s capabilities through simulations reflecting plausible biological and technical parameters of single-cell data, including cell sampling rate, phylogenetic reconstruction fidelity, cellular division and death rate, and show that PATH reproducibly and accurately measures heritability versus plasticity across different contexts. We show how the detection of heritability depends on sampling and phylogenetic reconstruction fidelity, and how these results can guide future lineage tracing experimental design and methods development. PATH can infer cell type transition dynamics with high accuracy, comparable to a classic maximum likelihood approach from species evolution [Lewis, 2001, Louca and Pennell, 2019, Pagel, 1994], but with higher computational efficiency, a critical feature considering the massive potential scale of phenotypically annotated phylogenies in high throughput single-cell data. We then apply PATH to published single-cell multi-omic datasets, which use either native or artificial lineage tracing (for human and model organism data, respectively), to explore two broad themes, development and cancer. Specifically, we examine mouse embryogenesis [Chan et al., 2019] and zebrafish neural development [Raj et al., 2018], a model of pancreatic cancer [Simeonov et al., 2021] and human glioblastoma [Chaligne et al., 2021]. PATH quantitatively maps cell fate trajectories during development, characterizes the variable plasticity of transcriptional states along the epithelial-to-mesenchymal transition in cancer and quantifies the heritability and stability of cell states of the corrupted neurodevelopmental hierarchy in glioblastoma. Finally, we apply PATH to newly generated single-cell whole genome sequencing data from a patient B-cell acute lymphoblastic leukemia (B-ALL) sample with a phylogeny constructed from somatic mutations with accompanying protein marker expression data. PATH reveals heritability of cellular phenotypes, and quantifies plasticity of immunotherapy-targeted B-cell surface markers and calculates transition rates between CD19 low, medium and high cell states. We make PATH available to the community as a comprehensive package, including software, analyses, and tutorials at https://github.com/landau-lab/PATH.

## Results

### Heritability, plasticity and cell state transition dynamics

Evolutionary biology offers a collection of metrics for characterizing heritable patterns of phenotypic variation, which can be adapted to interrogate single-cell ancestries. The degree to which phenotypic and ancestral similarity align is quantified by *heritability* statistics (*h*^2^ and *H*^2^) [Gillespie, 2004], which are weighted measures of the phenotypic correlation between relatives. These statistics have found application in agriculture, as part of the breeder’s equation, enabling the prediction of a phenotypic response to an artificial selection pressure [Gillespie, 2004]. Analogously, through leveraging phylogenetic trees, the degree to which related species phenotypically resemble each other, termed *phylogenetic signal* [Blomberg and Garland, 2002], can be quantified with various metrics (*e.g*., Pagel’s *λ* [Househam et al., 2022, Pagel, 1999], Blomberg’s *K* [Blomberg et al., 2003], Moran’s *I* [Gittleman and Kot, 1990]), and is used to make inferences about inheritance patterns and the evolutionary lability of phenotypes. These metrics are sometimes categorized as either statistic-or model-based [Münkemüller et al., 2012], but nonetheless show strong agreement [Diniz-Filho et al., 2012]. Signal statistics, such as Moran’s *I*, quantify the phylogenetic dependency of a phenotype, whereas model-based metrics, such as Pagel’s *λ*, assess the divergence between a phenotype’s phylogenetic distribution with a distribution expected by a model of random genetic drift. PATH builds upon these approaches to characterize the heritability or plasticity of cellular states in somatic evolution.

Specifically, PATH adapts Moran’s *I* (**Methods: Phylogenetic correlations**), a measure of *phylogenetic autocorrelation* and phylogenetic signal (but originally conceived as a spatial auto-correlation metric [Moran, 1950]), to quantify the heritability or plasticity of single-cell phenotypes. Like classic heritability statistics, phylogenetic autocorrelation is a measure of phenotypic similarity, weighted by relatedness. Phylogenetic auto-correlation quantifies the phylogenetic dependency of a single-cell measurement or phenotype (broadly defined), such as cellular state, transcriptional profile, or spatial location. Fundamentally, phylogenetic auto-correlation measures how much phenotypic resemblance close relatives have to one another compared to randomly chosen cells. If cells resemble close relatives much more than randomly chosen cells, the phenotype will appear highly heritable and phylogenetically auto-correlated. Such a pattern might be observed for a genetically encoded phenotype, as for example a phenotype affected by chromosomal copy number change. Alternatively, if closely related cells resemble each other to the same degree as any other cells, regardless of ancestry, the phenotype will appear plastic, not heritable and not auto-correlated. Such a pattern could reflect temporally transient states such as cell-cycle phase. Generally, phylogenetic auto-correlation captures the temporal stability or transience of a cell state, whether state is defined by intrinsic (*e.g*., mutation) or by extrinsic factors (*e.g*., interactions with the microenvironment). For example, if there is rapid toggling between states within a single generation, these states likely will not be auto-correlated in phylogenetic space, in contrast to more stable cell states that persist without transitioning for time scales longer than one cell division. Furthermore, we can assess statistical significance by computing phylogenetic correlation z scores, either analytically [Czaplewski and Reich, 1993] or by using a leaf-permutation test (**Methods: Phylogenetic correlations**). By measuring phylogenetic auto-correlations, PATH provides a powerful framework for quantifying the temporal stability and thus heritability versus plasticity of somatic cell states (or phenotypes) using multi-omic platforms that jointly capture the lineage history and the cell state of single cells.

In addition to quantifying the lineage dependency of single cell states to define heritability versus plasticity, to under-stand the evolutionary relationships *between* cell states we measure *phylogenetic cross-correlations* (**Methods: Phy-logenetic correlations**). Phylogenetic cross-correlation quantifies the dependency of one cell state’s distribution on the lineage patterning of another state. For example, again consider the phylogenetic distribution of a phenotype that depends on chromosomal copy number. If a chromosomal duplication occurs, cells with the extra chromosome, and affected phenotype, will be in close phylogenetic proximity to each other, and farther from cells without the chromosomal duplication. As such, each of the phenotypes, one affected and one unaffected by the duplication, will be autocorrelated, but because these phenotypes will be phylogenetically segregated from each other they will be negatively cross-correlated. On the other hand, if distinct measure-ments co-cluster phylogenetically, such as the transcription levels of two genes located on a chromosomal copy variant, such measurements will be positively cross-correlated. The phylogenetic cross-correlation of a cell state with itself is also its auto-correlation, so to simplify terminology when possible, we refer to both phylogenetic auto- and crosscorrelations as *phylogenetic correlations*.

To illustrate PATH, **Figure 1** depicts phylogenies that are the result of simulations of somatic evolution (**Methods: Simulating phylogenies**), where cells can transition between states. When cell states are heritable, meaning that state transitions occur infrequently (**Fig. 1A**), cells appear to phylogenetically group by state (*e.g*., **Fig. 1B**), and thus states are positively auto-correlated and negatively cross-correlated (**Fig. 1C,D**). In contrast, for highly plastic dynamics where state transitions occur frequently (**Fig. 1E**), cells do not appear to phylogenetically group by state (*e.g*., **Fig. 1F**), and states are lowly phylogenetically auto- and cross-correlated (**Fig. 1G,H**). The phylogenetic correlations between states can reflect evolutionary relationships; phylogenetic correlations increase or decrease with between-state transitions rates. For example, since transitions between state *α* and *γ* occur more frequently than transitions to *β* (**Fig. 1I**), *α* and *γ* co-cluster on the phylogeny (**Fig. 1J**) and are more phylogenetically correlated with each other than with *β* (**Fig. 1K,L**). Note that despite focusing on categorical cell states in **Figure 1**, phylogenetic correlations can also be computed for quantitative phenotypes (*e.g*., gene expression level).

**Figure 1:**
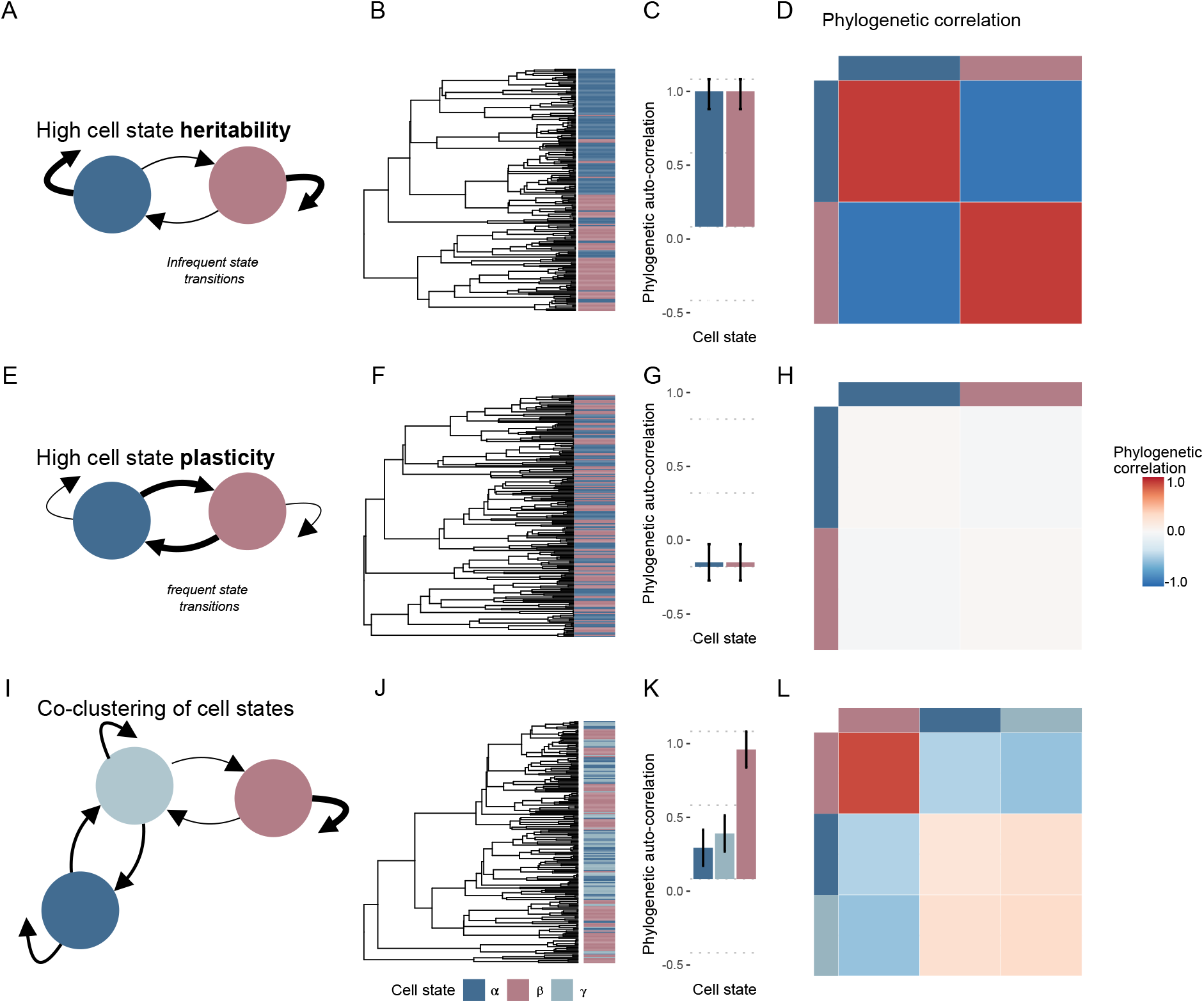
Phylogenetic correlations quantify the heritability versus plasticity of single-cell phenotypes. **A**) Diagram of highly heritable (categorical) cell state transition dynamics (**Methods: Markov model of cell state transitions**). Markov transition probabilities between states were simulated as *P_αα_* = *P_ββ_* = 0.9, and *P_αβ_* = *P_βα_* = 0.1 (meaning that cells had a 10% probability of switching states over each time point). **B**) Phylogenetic tree containing 200 cells, simulated as a somatic evolutionary process (**Methods: Simulating phylogenies**), from simulated transition dynamics depicted in **A**, with birth rate = 1 and death rate = 0. **C**) Phylogenetic auto-correlations (**Methods: Phylogenetic correlations**) for cell states depicted in **B**. **D**) Phylogenetic cross-correlation (**Methods: Phylogenetic correlations**) heat map for cell states depicted in **B**. Diagonals are equivalent to bars shown in **C**. **E**) Diagram of highly plastic (categorical) cell state transition dynamics (**Methods: Markov model of cell state transitions**). Markov transition probabilities between states were all the same (*P_αα_* = *P_ββ_* = *P_αβ_* = *P_βα_* = 0.5; meaning that cells had a 50% probability of switching states at any time). **F**) Phylogenetic tree containing 200 cells, simulated as a somatic evolutionary process (**Methods: Simulating phylogenies**), from simulated transition dynamics depicted in **E**, with birth rate = 1 and death rate = 0. **G**) Phylogenetic auto-correlations (**Methods: Phylogenetic correlations**) for cell states depicted in **E**. **H**) Phylogenetic cross-correlation (**Methods: Phylogenetic correlations**) heat map for cell states depicted in **F**. **I**) Diagram of a three-state system (**Methods: Markov model of cell state transitions**) in which states *α* and *γ* transition to each other at a rate higher than either transitions to state *β*. Markov transition probabilities between the three states were *P_αα_* = *P_αγ_* = *P_γγ_* = 0.5, *P_αβ_* = *P_βα_* = 0, *P_γα_* = 0.45, *P_βγ_* = 0.1, *P_γβ_* = 0.05, and *P_ββ_* = 0.9. **J**) Phylogenetic tree containing 200 cells, simulated as a somatic evolutionary process (**Methods: Simulating phylogenies**), from simulated transition dynamics depicted in **I**, with birth rate = 1 and death rate = 0. **K**) Phylogenetic auto-correlations for cell states depicted in **J**. **L**) Phylogenetic cross-correlation (**Methods: Phylogenetic correlations**) heat map for cell states depicted in **J**. Error bars in **C**, **G**, and **K** represent the analytical phylogenetic auto-correlation standard deviations calculated with the method from Czaplewski and Reich [1993].

We hypothesized that as cell state phylogenetic patterning can be related to the rate of state transitions (as in **Figure 1**), the rates of these state transitions might be inferred from such patterns. To test this, we simulated categorical state transition dynamics on idealized phylogenies (*i.e*., completely sampled and balanced, where every node has the same number of progeny; **Methods: Simulating phylogenies**, **Fig. S1A**). First, we confirmed a strong association between simulated transition rates and phylogenetic correlations (**Fig. S1B,** Spearman’s *ρ* = 0.89). Next, we explicitly connected phylogenetic correlations with a mathematical model of state transition rates (**Methods: Phylogenetic correlations and cell state transitions**, **Box S1**). For categorical cell states, phylogenetic correlations characterize the frequencies at which states are found within cell pairs that share recent ancestry, and these frequencies can be anticipated given a model of state transitions. For example, the states found within a pair of sister cells will depend on the state of the sisters’ shared parent and the rates at which transitions to other states can occur. For a highly heritable cell state in which transitions to other states occur infrequently, we will observe more sister cell pairs in the same such state than what we would expect given the state’s frequency. Using this mathematical relationship we can transform phylogenetic correlations into transition rate estimates with high accuracy (**Methods: Inferring cell state transitions from phylogenetic correlations**, **Fig.S1C**, **Box S1**).

### Measuring heritability, plasticity, and cell state transition dynamics in somatic evolution

The study of somatic evolution requires addressing an array of complicating biological and technical features not represented by idealized phylogenies (*e.g*., **Fig. S1A**). For instance, when cell division is not synchronized within a population [Brody et al., 2018], meaning that different cell generations coexist, the resultant phylogenies will be more adequately modeled in continuous-time. Additionally, not all cells will leave the same number of progeny, resulting in less balanced phylogenies. Moreover, in experimental contexts, not all cells are successfully assayed, leading to incomplete sampling. Other technical factors, such as sequencing depth or barcode length, can limit the detection or accumulation of heritable markers necessary to resolve close phylogenetic relationships. As such, to test the robustness of PATH across a wide range of biological and technical factors, we applied PATH to phylogenies simulated with a more sophisticated model of somatic evolution [Louca, 2020, Nee et al., 1994] (**Methods: Simulating phylogenies**). In this model, cell division and death occur, each with some probability, until the population reaches a chosen size. Then only a fraction of surviving cells is sampled and lineage relationships recovered. Cell states are simulated along the sampled phylogenies using a Markov model (**Methods: Markov model of cell state transitions**). Cell division, death, sampling, and state transition rates can be specified, thus providing a more accurate representation of somatic evolution to assess PATH’s applicability to complex somatic evolution datasets.

Consistent with our observations on idealized phylogenies (**Figure S1**), in phylogenies produced by this sampled somatic evolutionary process, phylogenetic correlations remain strongly related to cell state transitions. For instance, auto-correlation, our measure of heritability, declines as state transitions become more frequent. However, in addition to declining with plasticity, phylogenetic auto-correlations also decrease as sampling becomes sparser (**Fig. 2A**), underestimating heritability. Here, heritability is underestimated because incomplete sampling leads to an overestimation of lineage proximity in terms of node distance (**Fig. 2B**). In other words, cells that may appear to be close relatives on the tree (*e.g*., separated by one node) may in fact be more distant relatives due to the loss of unsampled intermediates (due to cell death, incomplete sampling or incomplete phylogenetic reconstruction). As such, when sampling is low, as might be the case when only hundreds or thousands of cells from a tumor are collected, even the closest related sampled cells from such lineages will usually represent fairly distant relationships, thus affecting heritability estimates. In these cases, only highly heritable phenotypes, reliably propagated over the number of cell divisions separating the closest related sampled cells will be detectable. These data reveal that under sufficiently sparse sampling, heritable phenotypes may appear plastic.

**Figure 2:**
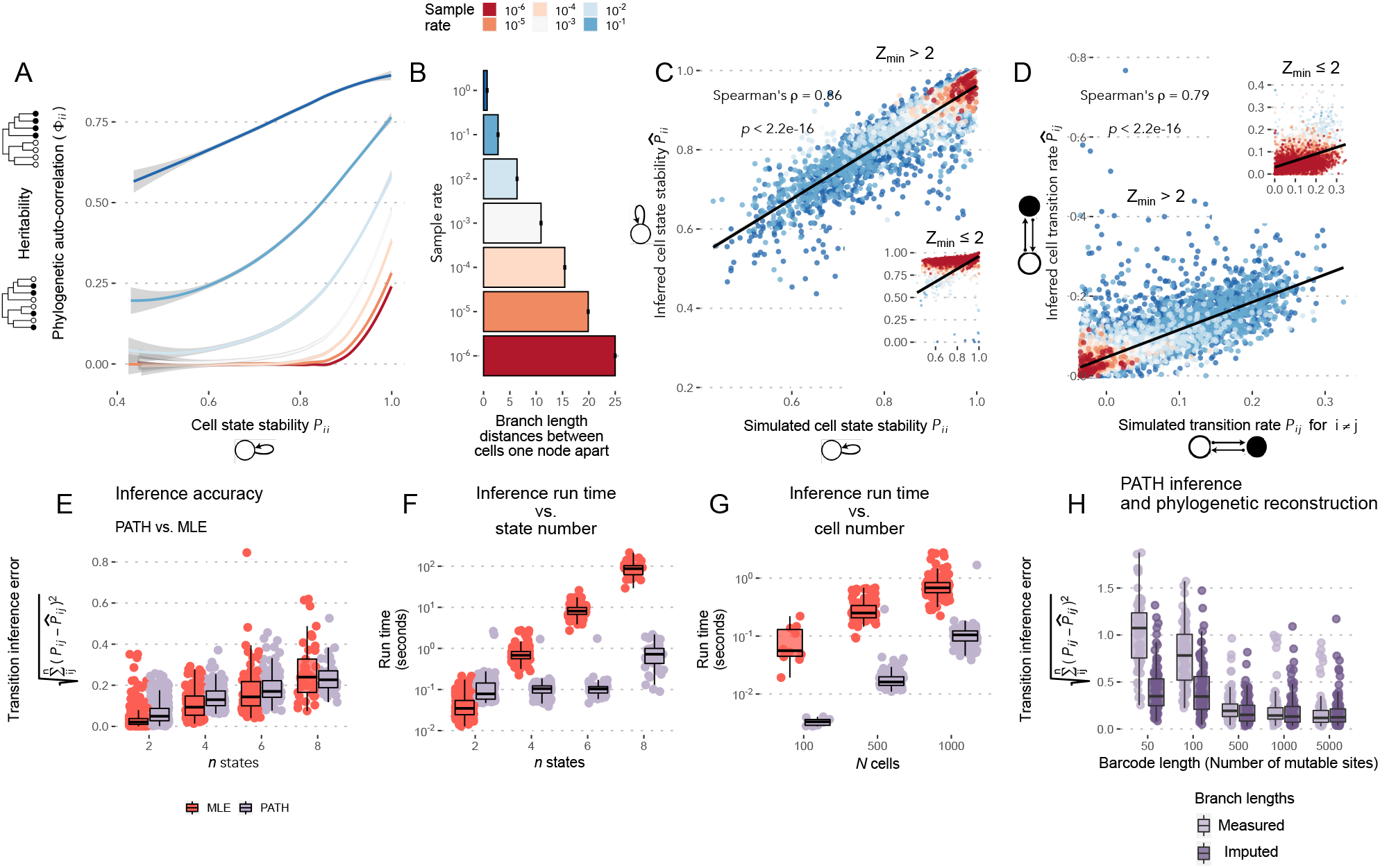
Measuring heritability, plasticity, and cell state transition dynamics in somatic evolution. **A**) Simulated cell state stability (Markov self-transition probability, **Methods: Markov model of cell state transitions**) for state 1 versus measured phylogenetic auto-correlation under different sampling rates (**Methods: Phylogenetic correlations**). Phylogenies contain 1,000 cells and Markov cell state transition dynamics were randomly generated for three-state systems. Phylogenies simulated as a sampled somatic evolutionary process (**Methods: Simulating phylogenies**) with birth rate 1 and death rate 0. Lines colored by sampling rate depict LOESS regression lines with 95% confidence intervals (light gray). **B**) Mean branch length (in units of time) distance between cell pairs only one-node apart on phylogenies versus cell sampling rate for phylogeny simulations. **C**) Simulated cell state stability (Markov self-transition probability) for state 1 versus PATH-inferred state stability for systems with phylogenetic auto-correlation z scores > 2. Colors represent sampling rates. Inset shows systems with at least one phylogenetic auto-correlation z score *≤* 2, and uses the same regression line. **D**) Simulated versus PATH-inferred cell state transition probability from state 1 to state 2 for three-state systems with phylogenetic auto-correlation z scores > 2. Colors represent sampling rates. Inset shows systems with at least one phylogenetic auto-correlation z score *≤* 2, and uses the same regression line. **E**) Comparing the state transition dynamic inference accuracy of PATH (light purple) with Maximum Likelihood Estimation (MLE; orange). Inference error is calculated as the Euclidean distance between inferred and simulated transition probability matrices (equation shown on y-axis label), and the number of possible states in a simulated system is shown on the x-axis (**Methods: Assessing cell state transition inference accuracy**). Panel depicts simulations for 1,000 cell phylogenies, sampled at a rate of 10^-2^, excluding simulations in which either inference method failed (which were usually due to the complete absence of some cell states). **F**) Same as **E** but measuring compute time. **G**) Comparing PATH and MLE compute times while varying phylogenetic tree size (number of cells; x-axis) fixing systems to four cell states, and sampled at 10^-2^. All inferences filtered to simulations surpassing the minimum phylogenetic auto-correlation z score threshold of 2. **H**) Comparing state transition inference of PATH using two different node depth estimation methods: (light purple) using measured branch length distances, and (dark purple) using imputed branch lengths (**Methods: Imputing branch lengths**) from estimated cell sampling rates. Simulations are for three-state systems simulated on 1,000 cell sampled somatic evolutionary phylogenies (**Methods: Simulating phylogenies**). Phylogenies were reconstructed by using the UPGMA algorithm on the cell pairwise Hamming distances between simulated lineage barcodes that were stochastically scarred at rate *s* = 0.01 (**Methods: Phylogenetic reconstruction**).

Next, we used PATH to infer state transition dynamics on phylogenies simulated by the sampled somatic evolutionary process. Since our inference approach transforms heritability measurements – which are underestimated when sampling is low – into transition rate estimates, transition inference accuracy was highest when state heritabilities were detectable (state auto-correlation z scores > 2, **Fig. 2C,D,** insets depict inferences for simulations in which heritability was not detectable [z score *≤* 2]). Notably, transition inference accuracy (**Methods: Assessing cell state transition inference accuracy**) with PATH is comparable to state-of-the-art Maximum Likelihood Estimation (MLE) methods (as implemented in Louca and Doebeli [2018]) traditionally used in evolutionary biology to infer character transitions (**Fig. 2E**, **Fig. S2A,B**), but with significantly faster compute times when analyzing a large number of states (**Fig. 2F**, **Fig. S2C**) and/or cells (**Fig. 2G**, **Fig. S2C**). PATH’s relative speed derives from the fact that PATH transforms a statistic (phylogenetic correlation) into a transition probability, whereas MLE uses an optimization algorithm to search for the most likely state transition probabilities and often requires many more calculations.

Another important confounder in harnessing phylogenetic trees to measure heritability is the fidelity of phylogenetic reconstruction. Intuitively, this can be understood in the context of artificial lineage tracing techniques that stochastically scar or cut genetic barcodes (*e.g*., Molecular recorder [Chan et al., 2019] and scGESTALT [Raj et al., 2018], where a limited number of cut sites can result in phylogenetic reconstruction errors. To understand this, beyond simulating phylogenies as a sampled somatic evolutionary process, we also simulated the reconstruction of these phylogenies by employing a model of CRISPR/Cas9 scarring (**Methods: Phylogenetic reconstruction**). To do this, each cell in a simulated evolving population contains a *barcode*, or a set of mutable and heritable sites that can be modified (*i.e*., scarred) stochastically. In contrast to our previous approach in which true phylogenies were recovered, here phylogenies were reconstructed from the differences between barcodes retrieved from cells in the terminal population, much as they would be for lineage tracing experiments. Comparing reconstructed with true phylogenies, we observe that as the number of mutable sites or barcode length increases, phylogenetic reconstruction accuracy improves (**Fig. S2D**). Concordant with reconstruction accuracy, state transition inferences using PATH also improve (**Fig. 2H**).

Since the accuracy of state transition inferences using PATH is affected by reconstructed branch lengths, which scale phylogenetic distances by time, inference will be impeded when branch lengths are inaccurate, and not possible when branch lengths are absent (which is common for single-cell phylogenies using artificial scarring methods). PATH can compensate for this by imputing terminal branch lengths, independent of phylogenies, if cell population sizes can be approximated (**Methods: Inferring cell state transitions from phylogenetic correlations, Imputing branch lengths**). PATH achieves this because under the model of sampled somatic evolution, the degree by which sampling leads to an overestimate of phylogenetic proximity can be calculated (**Fig. 2B**, **Fig. S2E,F**) and accommodated. In other words, under incomplete sampling, in which close phylogenetic relationships are overestimated due to the loss of unsampled intermediate cells, from the sampling rate (and independent of the reconstructed phylogeny), we can estimate how many intermediates are unsampled, and rescale branch lengths accordingly. Replacing measured branch lengths with model-imputed lengths significantly improves the accuracy of state transition inferences using PATH, particularly for low fidelity phylogenetic reconstructions where branch lengths are often less accurate (**Fig. 2H**). Thus, using PATH, state transitions can be accurately inferred for low fidelity phylogenies and when branch lengths are absent (in contrast to MLE), making PATH a powerful tool for the analysis of phylogenies produced by molecular scarring technologies.

In conclusion, these simulated datasets demonstrate that PATH, through the measurement of phylogenetic correlations, provides a comprehensive framework to analyze cell state heritability and plasticity in somatic cell populations, and can transform these measurements into inferences of state transition dynamics. PATH can accommodate a wide range of biological and technical features associated with somatic evolution. Thus, observable patterns of heritability and plasticity are robustly linked to the (often unobservable) processes that produce them, providing insights into cell lineage histories and somatic evolutionary dynamics. Having explored PATH’s capabilities on simulated datasets, we next sought to apply PATH to published single-cell lineage tracing datasets in two broad contexts, development and cancer.

### PATH quantifies ancestry and divergence of germ layers and cell types during mouse embryogenesis

Embryogenesis and organogenesis require the organization of the progeny of progenitor cells, which are restricted in number, location and levels of potency, into complex tissues. Single-cell lineage tracing methods provide sufficient resolution to map the cellular trajectories and interactions that underlie this exquisitely regulated organization. We reasoned that the application of PATH to such datasets would enable quantification of cell differentiation patterns through calculation of (i) phylogenetic auto-correlations that can be interpreted in this developmental context as cell state commitment strength and (ii) phylogenetic cross-correlations to determine relationships between tissue layers and cell types, and to understand gene expression across development.

We first asked whether PATH is able to reconstruct known cell fate relationships and dynamics in the well-characterized context of murine gastrulation (**Fig. 3A**). To accomplish this, we applied PATH to published mouse embryogenesis data [Chan et al., 2019], comprising single-cell phylogenies with matching single-cell transcriptional data. The authors leveraged a CRISPR/Cas9 lineage tracing construct to study early murine development, isolating embryos at E8.5 and constructing phylogenies from the edited barcodes (**Fig. 3B, Fig. S3A**). We applied PATH to these data to measure phylogenetic correlations for cellular phenotypes at multiple levels of resolution, and gained insight into the commitment and divergence patterns of cellular phenotypes from their origin layers in the blastocyst through gastrulation, and ultimately to their differentiated tissue in the E8.5 embryo.

**Figure 3:**
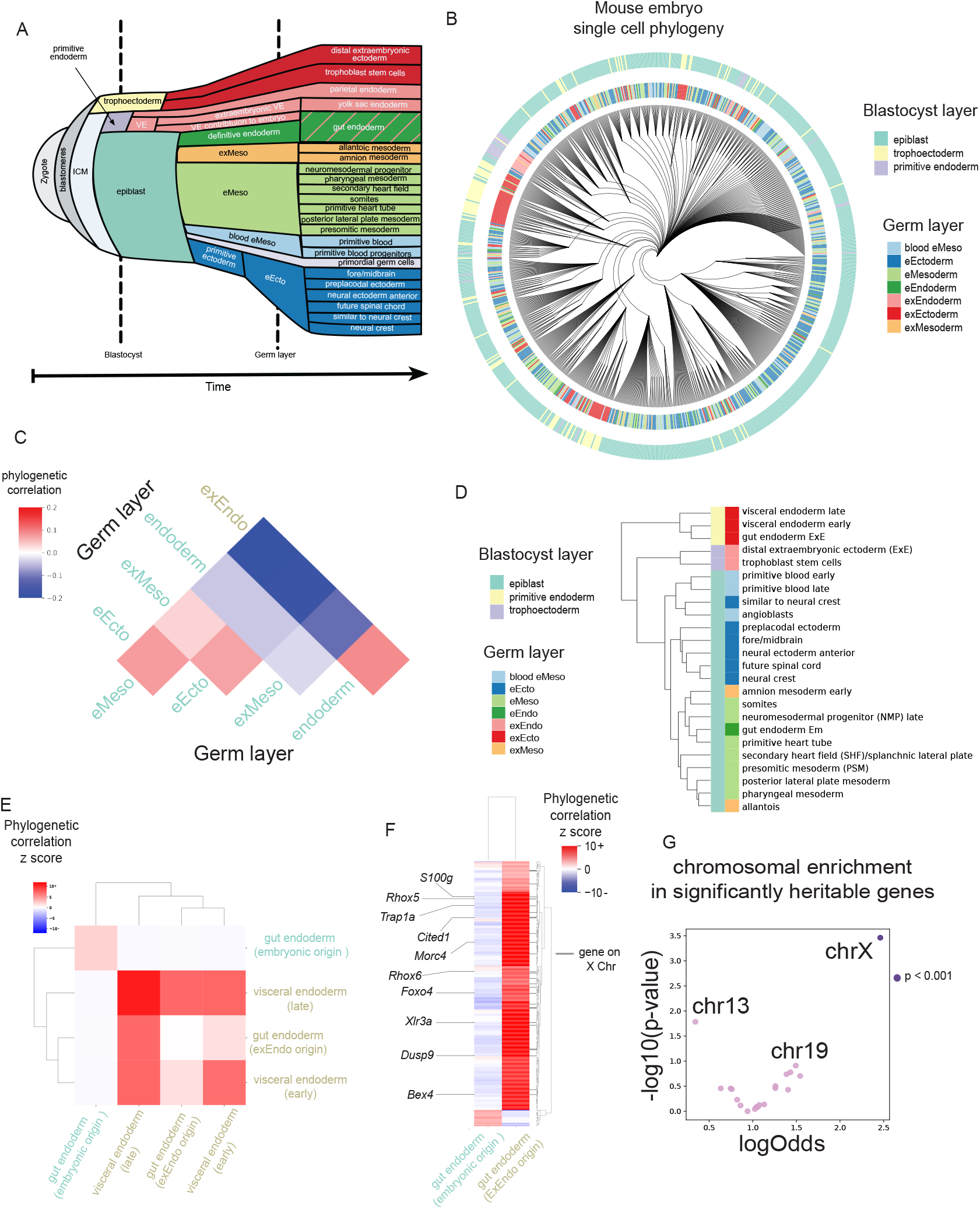
PATH quantifies ancestry and divergence of germ layers and cell types during mouse embryogenesis. **A**) Schematic of mouse embryogenesis adapted from Thowfeequ and Srinivas [2022]. VE, visceral endoderm; ICM, inner cell mass; e prefix, embryonic; ex prefix, extraembryonic. **B**) Single-cell phylogeny from mouse embryo 6 from Chan et al. [2019], containing 700 randomly chosen of 1,722 cells for visualization. Each leaf represents a single cell. Leaves are colored by blastocyst or germ layer of origin. e prefix, embryonic; ex prefix, extraembryonic. **C**) Germ layer phylogenetic correlations for embryo 2. Labels colored by cell type blastocyst origin: visceral endoderm, gold; epiblast, green. **D**) Hierarchical clustering of tissue types by phylogenetic correlation using Ward’s method. Only tissues with more than 30 cells were used. Tissues colored by germ and blastocyst layer of origin. Phylogenetic correlations can be found in **Fig. S3C**. ExE, extraembryonic; EM, embryonic. **E**) Phylogenetic correlation z score of gut endoderm cells annotated by their source tissue in the blastocyst and visceral endoderm (early and late). Labels colored by cell type blastocyst origin: visceral endoderm, gold; epiblast, green. **F**) Phylogenetic correlation z scores between genes and tissue assignment. Genes on the X chromosome are denoted with a gray bar (right) with select X-chromosome genes labeled (left). Cell state labels colored by cell type blastocyst origin: visceral endoderm, gold; epiblast, green. The complete set of phylogenetic correlations are in **Table S1**. **G**) Enrichment of highly heritable genes at the whole chromosome level (with chromosome 13, 19 and X labeled). Log odds ratio and p-value (p < 10^-3^, Fisher’s exact test) of number of highly heritable genes (z score > 3) on each chromosome compared to all other chromosomes Only expressed genes were considered for comparison (top 2,000 most variable genes across phylogeny, see **Methods: Mouse embryogenesis**).

As expected, all blastocyst layers with sufficient representation had high auto-correlation in both replicates, indicating that a cell from a particular blastocyst layer is more likely to produce progeny that are also found in the same layer, reinforcing what is known about the rigidity of developmental programs [Thowfeequ and Srinivas, 2022]. Germ layers derived from outside of the epiblast had high auto-correlation in all replicates that had sufficient cell recovery, while tissues that shared a common origin in the epiblast had lower autocorrelations (**Fig. S3B)**. Thus, the non-epiblast-derived layers show evidence of earlier fate commitment, while the more plastic phenotype of the epiblast is consistent with its later divergence [Thowfeequ and Srinivas, 2022]. PATH also accurately reconstructed the patterns of shared ancestry between blastocyst layers and germ layers (**Fig. 3C**). Notably, phylogenetic correlations recovered the dual contribution of both embryonic- and extraembryonic-derived tissues to the endoderm [Kwon et al., 2008, Nowotschin et al., 2019, Pijuan-Sala et al., 2019] (**Fig. 3C**). This highlights PATH’s ability, by leveraging phylogenies, to identify phenotypically similar but ancestrally distinct cells.

After implementing PATH at the level of the blastocyst and germ layers, we sought to quantify the degree of shared origin of higher resolution, transcriptionally defined cell types derived from each germ layer (**Fig. 3D**). Cell types that share ancestry will likely be highly phylogenetically correlated. Indeed, PATH analysis correctly identified important developmental relationships between primitive blood cells (early and late); and neural crest and future spinal cord. Interestingly, PATH also identified the shared origins of the embryonic splanchnic lateral plate and extraembryonic allantois cells in the nascent mesoderm [Thowfeequ and Srinivas, 2022], highlighting PATH’s ability to identify shared ancestry from progeny that have diverged into different germ layers (**Fig. S3C,D**). Of note, we again observed high cross-correlation between the endoderm and extraembryonic endoderm-derived tissues in the gut endoderm (**Fig. 3C**), now at the level of cell type (**Fig. 3E**). This higher resolution analysis revealed that extraembryonic-derived endoderm tissue cross-correlates almost exclusively with cells from the late visceral endoderm (arising around E8.0 in the extraembryonic endoderm), as opposed to the early visceral endoderm (arising around E7.0 in the extraembryonic endoderm) [Grosswendt et al., 2020] or embryonic-derived gut endoderm. Given that the intercalation of extraembryonic endoderm into the gut endoderm occurs between E7.5 and E8.5 [Nowotschin et al., 2019], this analysis nominates a specific cell population from the extraembryonic visceral endoderm contributing to the definitive endoderm.

Having examined the phylogenetic correlations of embryonic germ layers and cell types, we then took advantage of the versatility of PATH to evaluate the heritability of gene expression programs in these populations of endoderm cells. We calculated phylogenetic correlations between each population of endoderm cells (originating in the epiblast or the primitive endoderm) and gene expression across the tree. We found distinct gene expression profiles phylogenetically correlated with each population of endodermal cells (**Fig. 3F**). In concordance with prior work, we found that *Rhox5* and *Trap1a*, two X-linked genes, had high phylogenetic correlation with endoderm cells with extraembryonic origin [Nowotschin et al., 2019, Pijuan-Sala et al., 2019]. Interestingly, we found that genes on the X chromosome beyond *Trap1a* and *Rhox5* were significantly enriched in this heritable expression program (**Fig. 3F,G**). This signal is grounded in the differential imprinting patterns between extraembryonic and embryonic cells: extraembryonic endoderm cells have paternally imprinted X-inactivation [Takagi and Sasaki, 1975] imbuing them with a unique expression pattern that has been shown to persist after intercalation into the visceral endoderm [Loda et al., 2022]. These results demonstrate PATH’s ability to explore patterns and timing of coordinated gene expression during development, including epigenetically propagated signals.

### PATH identifies cell fate-determining factors across anatomical, defined tissue and gene expression layers during neurogenesis in zebrafish

One notable aspect of PATH is its ability to quantify relationships between different types of phenotypic information, providing the opportunity to leverage not only transcriptional information from scRNAseq data, but also any available spatial, anatomical or temporal information. As such, we can perform multi-modal analysis to characterize relationships between these phenotypic annotation layers, and thus draw inferences about their interactions (for example, we can use the phylogenetic cross-correlations of individual genes with either cell or tissue type to nominate cell fate determination factors). To explore this capability, we applied PATH to prospectively lineage-traced developing zebrafish brains [Raj et al., 2018]. The data in Raj et al. [2018] comprise cells annotated not only by single-cell transcriptional profiling but also by the anatomic region from which they were dissected. These multi-layer annotations enabled us to investigate neuronal development dynamics within, between and across anatomically distinct brain regions.

We first used PATH to examine phylogenetic correlations of different brain regions. Neuronal tissue had been collected from two whole brains and anatomic regions were manually separated during dissection, resulting in three main regions (forebrain, midbrain, hindbrain; **Fig. 4A,B**). By projecting anatomic region on the reconstructed phylogeny and applying PATH, we found that each defined anatomic location had high phylogenetic auto-correlation, indicating that neuronal cells within a brain region share recent ancestry (**Fig 4C**). As expected, the cells with ambiguous annotations (labeled “mix”) had much lower phylogenetic auto-correlations, most likely due to heterogeneous sampling that diluted the phylogenetic signal.

**Figure 4:**
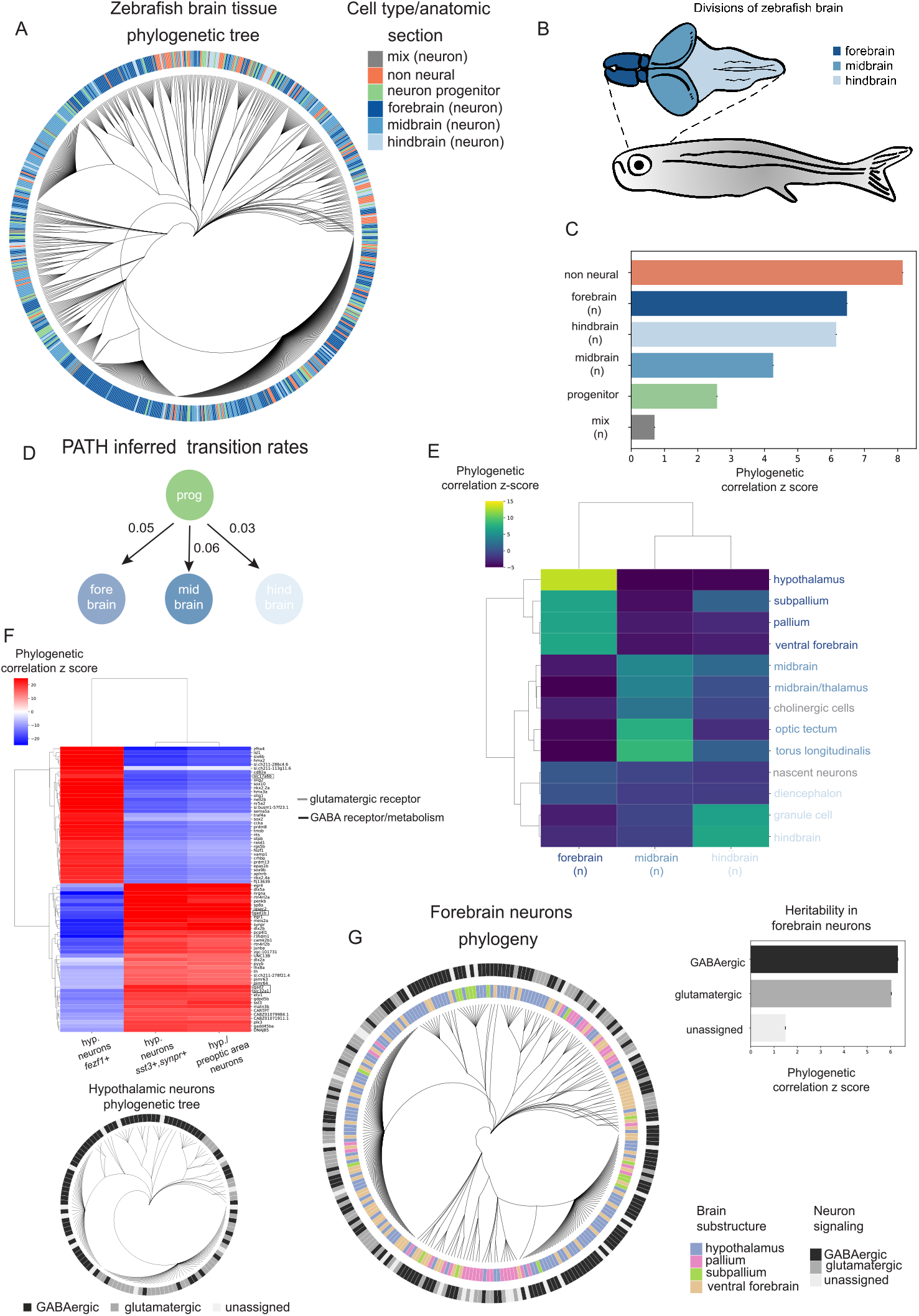
PATH identifies cell fate-determining factors across anatomical, defined tissue and gene expression layers during neurogenesis in zebrafish. **A**) Single-cell phylogeny from zebrafish brain 3 (replicate 1) from Raj et al. [2018]. Each leaf represents a single cell (N = 750). All cell type and anatomic section annotations are as defined in Raj et al. [2018], by scRNAseq and manual dissection, respectively. Cells colored in orange are non-neurons, cells in green are neural progenitors. Neuronal cells (blue hues and gray) are colored by the anatomic location from which they were dissected. Non-neural and neuron progenitor cells lack anatomical annotation. Cells labeled “mix” were from dissections with ambiguous anatomical origin (see **Methods: Zebrafish brain development**). **B**) Zebrafish brain schematic. Forebrain, midbrain and hindbrain have been labeled. **C**) Cell type/anatomic-section phylogenetic auto-correlations. Mature neurons are labeled “n” and annotated by dissection site (blues, gray); neuronal progenitors are labeled in green and non-neural cells are in orange. **D**) PATH inferred transition probabilities between neuron progenitor cells (prog) and neurons from each anatomic brain region. Branch lengths imputed by approximating the cell sampling rate to be 10^-4^ to infer transition probabilities. Values rounded to the nearest hundredth. **E**) Phylogenetic correlation z scores between anatomic site and transcriptionally assigned brain substructure across all neurons. Substructures are colored by brain location from **A**. **F**) Phylogenetic correlation z scores between (top 2,000 most variably) expressed genes and individual hypothalamus clusters (defined by Raj et al. [2018] from select marker genes). The 35 most auto-correlated genes per cluster are shown, and a complete set of phylogenetic correlations are in **Table S2**. Phylogenetic tree of hypothalamic neurons annotated by GABA/Glut signaling (**Fig. S4C**) (see **Methods: Zebrafish brain development**). **G**) (Left) phylogeny of all forebrain neurons (N = 270), leaves annotated by brain substructure assignment and GABA and glutamatergic signaling. (Right) phylogenetic auto-correlation of GABA and glutamatergic signaling across all forebrain neurons.

To characterize potential developmental trajectories between neurons and neuronal progenitors, we next used PATH to infer transition dynamics between them, segregating neurons by their anatomic region. Notably, we found that the progenitor cell pool contributes at similar rates to the forebrain, midbrain and hindbrain (**Fig. 4D**), consistent with the findings of Raj et al. [2018] suggesting that progenitor cells were multipotent at the time of barcoding.

As the versatility of PATH allows not only for comparisons within the same category of data (*e.g*., brain region), but also for integrated analysis across different layers of phenotypes, we next aimed to examine the phylogenetic correlation of anatomical brain regions with higher-resolution brain structure information derived from scRNAseq marker data. PATH analysis showed that these brain structures cross-correlate with their expected anatomical region (**Fig. 4E**), demonstrating the ability to correctly integrate transcriptionally and anatomically derived single-cell annotations across a phylogeny.

We next focused our analysis on the hypothalamus, a complex brain structure that is essential for the maintenance of homeostasis in an organism’s adaptive response to its environment. This structure is composed of a variety of anatomically and molecularly distinct neuron subtypes which respond to and release distinct sets of neuropeptides and hormones [Benevento et al., 2022]. Given this complexity, the transcriptional and phylogenetic dynamics underlying the functional organization of the hypothalamus were of interest for us to explore within the PATH framework. Using gene clusters defined by Raj et al. [2018] using scRNAseq, we first assessed the phylogenetic correlations of transcriptionally distinct clusters **(Fig. S4A**) of hypothalamic neurons. This analysis showed that *tac1*+, *nrgna*+, neurons were highly cross-correlated with neurons from the preoptic area (POA), indicating a shared cellular ancestry. The expression of both of these genes was negatively cross-correlated with *fezf1* + neurons, indicating distinct histories (**Fig. S4A**). To explore the molecular underpinnings of these differences in developmental origins we cross-correlated gene expression with hypothalamic neuron subtype (**Fig. S4A**) across the phylogeny of forebrain neurons to determine which genes were most strongly cross-correlated with these cell types (**Fig. 4F**). Interestingly, we found that genes required for glutamatergic signaling (*slc17a6b*) were highly cross-correlated with *fezf1*+ neurons, while those genes required for GABAergic signaling (*gad1b*, *gad2*, *slc32a1*) were highly cross-correlated with POA and *tac1*+, *nrgna*+, neurons, indicating that use of GABAergic or glutamatergic signaling is a heritable trait in cells of the differentiating hypothalamus (**Fig. 4F**). Indeed, we found that glutamatergic and GABAergic signaling were heritable in the forebrain (**Fig. 4G, Fig. S4B,C**), consistent with lineage tracing studies that found high heritability of GABAergic signaling in the murine forebrain [Bandler et al., 2021]. Thus, PATH is able to connect gene expression profiles to cell state through lineage information in an unbiased, quantitative manner, and uncovers the contribution of biologically meaningful cell populations underlying the observed patterns of heritability.

### Quantifying cell state transitions during metastasis

Malignant populations harbor significant cell state diversity and the characterization of their relative heritability and plasticity is currently a major goal of the cancer field [Bell et al., 2019, Fennell et al., 2022, Oren et al., 2021, Shaffer et al., 2020]. Tumor single-cell phylogenies provide a unique opportunity to distinguish between cancer cell state heritability versus plasticity. Cancer cell state diversity has been associated with critical disease aspects such as tumor growth [Neftel et al., 2019], treatment response [Fennell et al., 2022], and metastatic spread [Karras et al., 2022], emphasizing the need to define the heritability versus plasticity of cancer cell states. Notably, in comparison to primary tumors, in most contexts there is a lack of established, recurrent genetic drivers of metastasis [Rogiers et al., 2022]. Thus, other non-genetic factors likely play a major role in metastasis. We therefore applied PATH to correlate lineage dynamics with key non-genetic features, including location and cell state, of metastatic tumors. We re-analyzed data from a murine model of metastatic pancreatic cancer with inducible CRISPR/Cas9 based lineage recording and scRNAseq [Simeonov et al., 2021]. Metastatic tumors are thought to arise by the dissemination of a single or a small number of clones from the primary tumor [El-Kebir et al., 2018, Gundem et al., 2015, Hu et al., 2019, Navin et al., 2011, Turajlic et al., 2018]. By leveraging PATH’s ability to integrate data of different modalities, we tested this assumption by assessing the shared ancestry of metastatic tumor cells harvested from distinct anatomical sites: primary tumor (pancreas), lung metastatic tumor, liver metastatic tumor, peritoneal metastatic tumor, tumors forming at the site of the surgical lesion and circulating tumor cells (CTCs). Cellular tissues of origin were highly phylogenetically autocorrelated (**Fig. 5A,B**), consistent with the established model in which a small number of founder cells seed metastases, creating site-specific clonal bottlenecks. Importantly, the quantification provided by PATH allowed for direct comparison of harvest site-specific lineages, revealing patterns of clonal seeding in metastasis. For instance, surgical lesions (which formed on the peritoneal surgical incision site) and peritoneal metastases had negative phylogenetic correlation, (**Fig. S5A**) suggesting that they had distinct origins despite their physical proximity. As expected, CTCs, which may have many distinct clonal origins, had lower phylogenetic auto-correlation than solid tissues (**Fig. 5B**).

**Figure 5:**
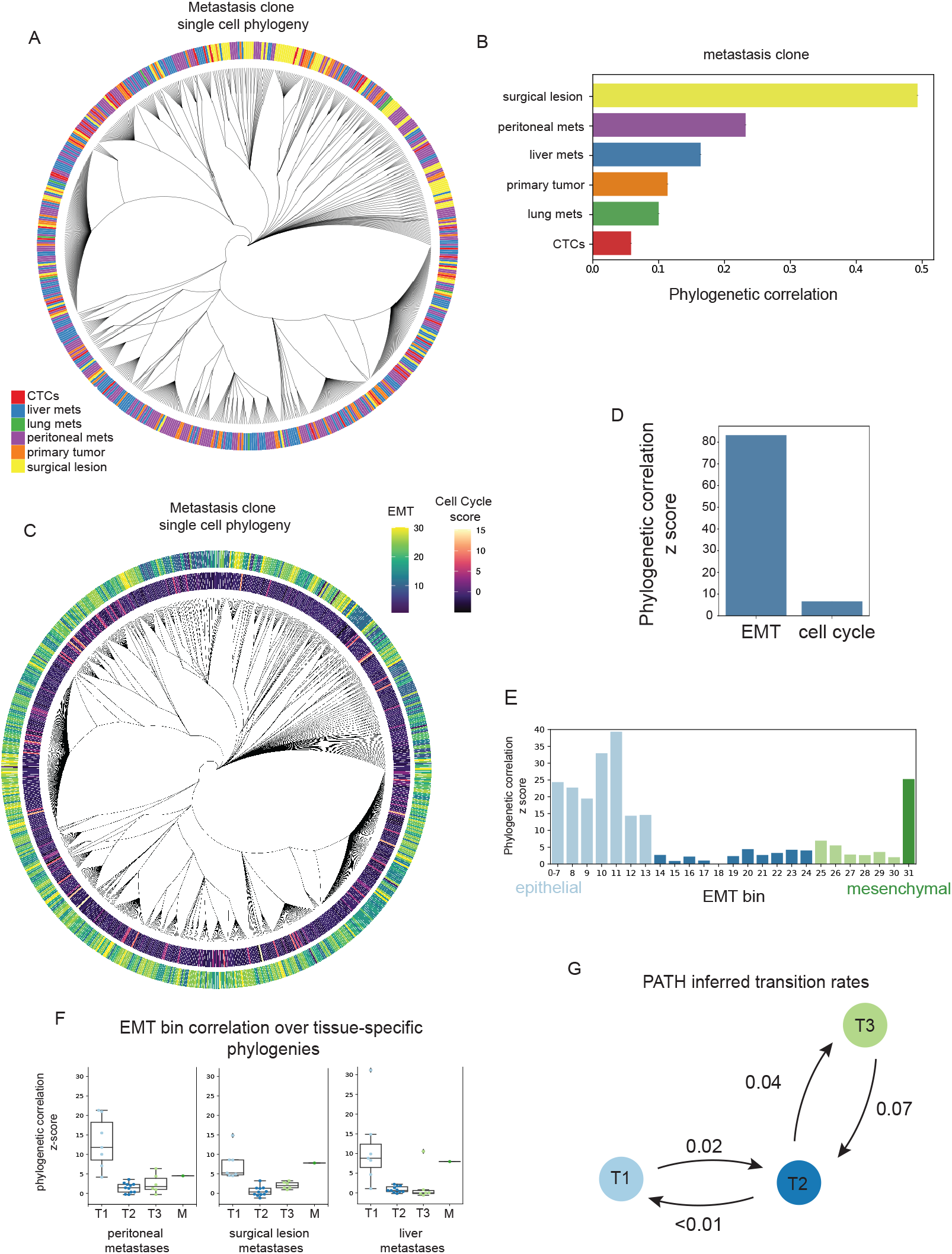
Quantifying cell state transitions during metastasis. **A**) Single-cell phylogeny from Mouse 1, Clone 1 from Simeonov et al. [2021], containing 700 randomly chosen of 7,968 cells for visualization. Each leaf rep)resents a single cell. Leaves are colored by their harvest site. CTCs denote circulating tumor cells. Mets, metastases. **B**) Phylogenetic auto-correlation of tumor cells annotated by harvest site. Bars colored by harvest site, as in **A**. **C**) Single-cell phylogeny from **A**, with cells colored by EMT and cell cycle score (G2M score). **D**) EMT and cell cycle phylogenetic auto-correlations across all tumor cells (N = 7,958). **E**) EMT bin phylogenetic auto-correlations (z scores) using all cells. Bins are colored by transition states derived from **Fig. S5B**. **F**) Box and whisker plot of EMT bin phylogenetic correlations (z scores) across phylogenies that contain cells from only one harvest site. Dots correspond to EMT bins. Bins are grouped and colored by transition state membership. Boxes represent the interquartile range (IQR); the center line represents the median; minima and maxima shown represent 1.5·IQR. **G**) PATH inferred transition probabilities between states (T1, T2, T3) using all cells (N = 7,968). Values rounded to the nearest hundredth. Transition probability inferences use imputed branch lengths by approximating a sampling rate of 10^-6^ (see **Methods: Mouse model of pancreatic cancer**).

The epithelial-to-mesenchymal transition (EMT) plays a crucial role in metastasis [Dongre and Weinberg, 2019, Lambert et al., 2017, Thiery, 2002], and thus Simeonov et al. [2021] calculated an EMT score for each tumor cell, reflective of that cell’s position along a transcriptional continuum from highly epithelial to mesenchymal cells. Low scores correspond to more epithelial characteristics and high scores correspond to more mesenchymal characteristics. Of note, there is an ongoing discussion in the field regarding whether EMT is best modeled as a series of functionally discrete, transcriptionally and epigenetically distinct intermediate states or a continuum of transcriptional hybrid states [McFaline-Figueroa et al., 2019, Pastushenko and Blanpain, 2019, van Dijk et al., 2018]. Because we can simultaneously observe both cellular position within the EMT continuum and on the phylogeny, this dataset offers a unique opportunity to investigate this question (**Fig. 5C**).

First, phylogenetic auto-correlation revealed the high heri-ability of cellular position on the EMT transcriptional continuum (**Fig 5D**). This finding can be contrasted with phylogenetic auto-correlation measurements of cellular position within the cell cycle, which can serve as a negative control, as position within the cell cycle is not usually expected to depend on ancestry [Chaligne et al., 2021] (**Fig 5C,D**).

Next, we asked how heritability and plasticity varied across the EMT continuum. Cells had been assigned EMT scores ranging from 0, denoting a completely epithelial cell to > 30 denoting a completely mesenchymal cell [Simeonov et al., 2021]. We partitioned cells along the continuum using units of 1 (bin #1 includes cells with EMT scores from 0 to 1, bin #2 includes cells from 1-2, *etc*.), merging bins at the extremes (all cells with a score of 7 or less were assigned to a single bin, as were cells that scored higher than 30) because these bins had low cellular representation. We calculated phylogenetic correlations for each individual bin, revealing four distinct groups of cross-correlated states along the EMT continuum defined by varying degrees of heritability (**Fig. 5E; Fig. S5B,C, Table S3**). Specifically, one group of phylogenetically correlated states corresponds to the epithelial and early transition states (T1), indicating that cells in this part of the EMT continuum tended to remain in the T1 state and were less likely to transition to other states. Likewise, mesenchymal (M) cells were also highly phylogenetically auto-correlated, indicating temporal stability of the mesenchymal state. However, cells in bins in the middle part of the continuum (later transition states; T2, T3) appeared less heritable, suggesting that these states were more plastic (**Fig. 5E, Fig. S5B**). These results were robust to different bin sizes (**Fig. S5D)**, suggesting that these results are not an artifact of the binning procedure. Intriguingly, these results imply that despite tumor cells occupying a continuum of EMT transcriptional states, the states at the extremes of the continuum show a higher degree of heritability, whereas intermediate cells states show a higher degree of plasticity. As our analysis above showed a high degree of phylogenetic similarity within the same metastatic location, we further ruled out that EMT heritability is driven by variability in the representation of EMT states across metastatic sites (**Fig. 5F**). Furthermore, these results were replicated within each metastatic location, and consistently showed the T1 state to be the most heritable within each tissue, and the T2/T3 states to be more plastic, suggesting that patterns of cell state heritability were not driven by tumor location.

Finally, to quantify cell state transitions from the initial epithelial state to the more plastic later states, we used PATH to infer transition dynamics between early (T1), middle (T2) and late (T3) EMT states. We observed that transitions out of the early epithelial state (T1) into more plastic states along the continuum (T2) occurred with some frequency, but transitions in the reverse direction going from a later plastic state back to an early epithelial state were rare. In contrast, we found marked plasticity between later intermediate states (T2 and T3) (**Fig. 5G**). These results suggest that EMT represents neither a smooth continuum of hybrid states nor an equally discretized cell state trajectory, but instead comprises punctuated states with different transition probabilities. These analyses indicate an integration of the two proposed models of EMT: cells undergoing EMT are transcriptionally continuous (as reported by [McFaline-Figueroa et al., 2019, Pastushenko and Blanpain, 2019, Simeonov et al., 2021, van Dijk et al., 2018]), but their lineage dynamics reveal functionally and heritably distinct states in EMT (as reported from functional transplantation assays in mice by Pastushenko et al. [2018]). These findings highlight the power of combining single-cell multi-omics data with phylogenetic information to draw conclusions that would not be possible through analyzing either data type alone.

### Elucidating heritable transcriptional modules and cell state transition dynamics in human glioblastoma

While artificial lineage tracing is a powerful approach in model organisms, it cannot be applied to reconstruct phylogenetic relationships in human data. Recent advances in multi-modal single-cell sequencing enable joint lineage reconstruction and cell phenotyping in primary human samples [Sankaran et al., 2022]. To examine this exciting frontier, we applied PATH to phenotypically annotated retrospective phylogenies reconstructed from human singlecell data leveraging stochastic DNA methylation changes as native lineage barcodes (**Methods: Human patient glioblastoma**) [Chaligne et al., 2021, Gaiti et al., 2019].

Having observed the high heritability of harvest site location across multiple tumors in metastasis (**Fig. 5A,B**), we set out to test whether a cell’s spatial location within a single tumor was stable. We applied PATH to MGH105, an IDH-wildtype (WT) glioblastoma (GBM) patient specimen in which cells were sampled from four distinct tumor locations (**Fig. 6A**) [Chaligne et al., 2021, Neftel et al., 2019]. We found that each of the locations (inset, **Fig. 6A**) were highly phylogenetically auto-correlated (leaf-permutation test, **Fig. 6B**), indicating that spatially proximal tumor cells were also more proximal in terms of ancestry, consistent with our expectations for a solid tumor malignancy.

**Figure 6:**
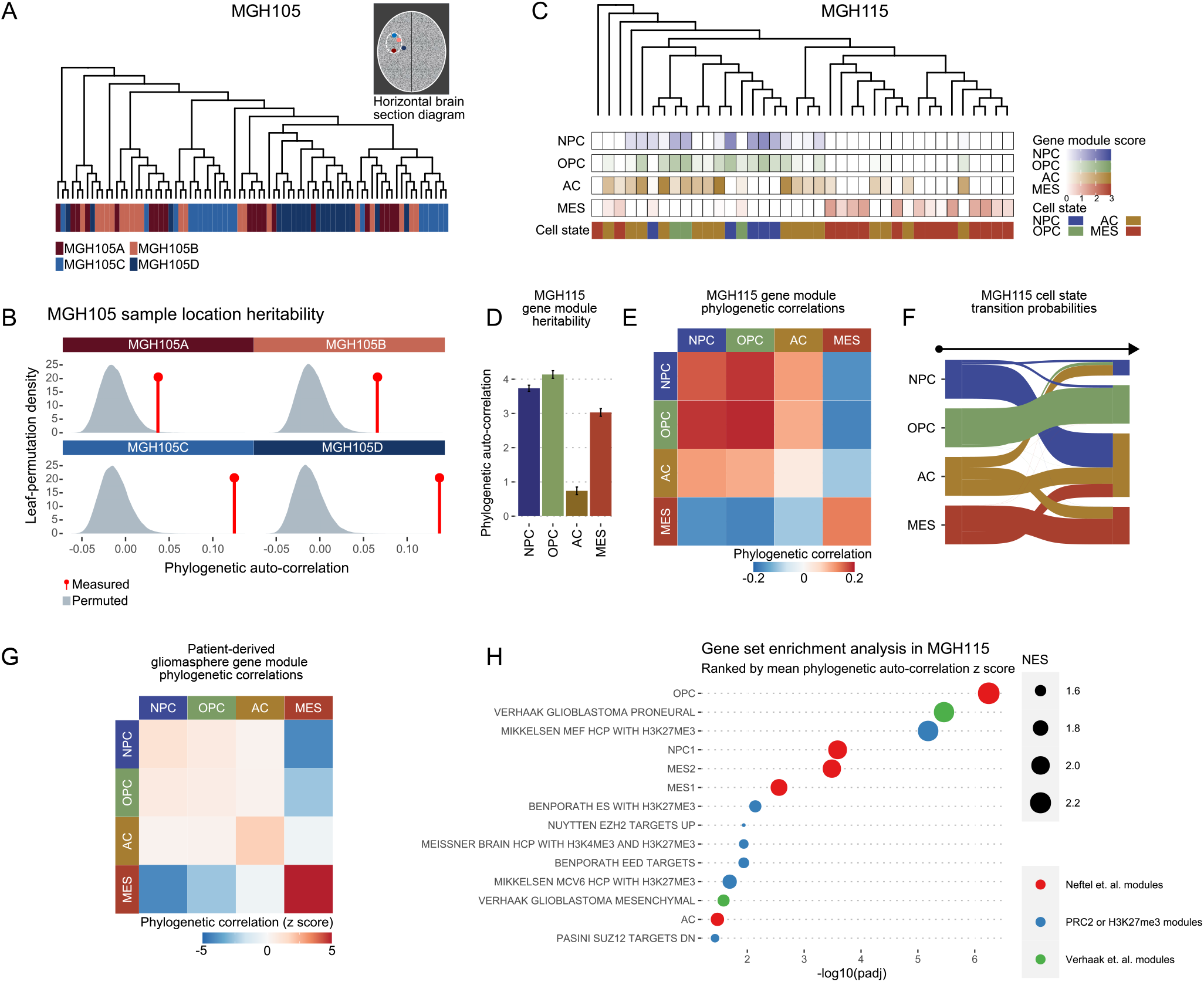
Heritable transcriptional modules and cell state transition dynamics in human glioblastoma. **A**) Human GBM sample (MGH105) single-cell consensus phylogeny containing 80 cells (20 from each tumor location) with tumor sample location projected onto leaves. Inset is a schematic of the four MGH105 patient tumor sample locations. **B**) Leaf-permutation test (10^6^ permutations) of tumor sample location phylogenetic auto-correlation. Density plot depicts leaf-permutation auto-correlations and red lines show measured (non-permuted) phylogenetic auto-correlations. **C**) Human GBM patient sample (MGH115) single-cell phylogeny (replicate 6) containing 38 cells with GBM gene module scores and categorical cell states projected onto leaves. **D**) Replicate mean (across 9 MGH115 phylogeny replicates) phylogenetic auto-correlation z scores for GBM gene module scores for patient sample MGH115. **E**) Replicate mean phylogenetic correlation heat map for patient sample MGH115 GBM gene modules. **F**) Sankey plot of replicate mean Markov transition probabilities inferred from categorical state phylogenetic correlations in patient sample MGH115 phylogeny replicates. Probabilities shown are shown for 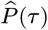 (**Methods: Inferring cell state transitions from phylogenetic correlations**). **G**) Replicate mean phylogenetic correlation z score heat map for gliomasphere GBM gene modules, using one-node weighting. **H**) Dot plot of enriched pathways from GSEA of chemical and gene perturbation curated gene sets (C2:CGP) and six GBM gene modules (NPC1-/NPC2-/OPC-/AC-/MES1-/MES2-like) [Neftel et al., 2019] for patient sample MGH115, with genes ranked by their phylogeny-replicate mean phylogenetic auto-correlation z scores (**Methods: Phylogenetic correlations**, **Methods: Human patient glioblastoma**). Only select gene sets are depicted; other significantly enriched gene sets can be found in **Table S5**. Dot sizes are proportional to GSEA normalized enrichment scores (NES). GBM gene modules (NPC-/OPC-/AC-/MES-like) were shortened to (NPC/OPC/AC/MES).

GBM harbors significant cell state diversity, which can be classified according to the expression four major gene modules, defined as neural progenitor-like (NPC-like), oligodendrocyte progenitor-like (OPC-like), astrocyte-like (AC-like), and mesenchymal-like (MES-like) [Neftel et al., 2019]. By measuring transcriptional signatures for these modules in each cell, GBM cells can be classified into four distinct transcriptionally-defined cell states. These cell states can be further grouped by function; for instance, we define the stem-like cells as cells that highly express one of the progenitor (NPC-or OPC-like) gene modules. The stem-like and AC-like states each resemble a known neurodevelopmental program, and thus can be collectively considered as neurodevelopmental-like. In contrast, the MES-like state does not reflect a developmental brain expression program and its emergence has been associated with both genetic and non-genetic factors, including interaction with immune cells and hypoxia [Hara et al., 2021].

The cell state heterogeneity in GBM has been a challenge for successful implementation of targeted therapies [Nicholson and Fine, 2021], so understanding the mechanisms and dynamics of cell state plasticity could provide insight into more effective treatment regimens. To examine the potential heritability or plasticity of these cell states, we re-analyzed MGH115, a human patient-derived GBM sample with annotated phylogeny with (i) continuous gene transcriptional module scores (generated from module-specific gene expression using matched scRNAseq) and (ii) categorical cellular state annotation based on the per cell maximum transcriptional module score (**Fig. 6C**). The stem-like (NPC-/OPC-like) and MES-like transcriptional modules displayed high phylogenetic auto-correlations, suggesting that in this specimen, the expression of these genes is in part heritable. The AC-like module, however, was not significantly phylogenetically auto-correlated, suggesting that the transcriptional state was more plastic in this patient sample (**Fig. 6D**).

As the MES-like state does not recapitulate any neurodevelopmental expression program and has been reported to be influenced by non-genetic factors [Hara et al., 2021, Neftel et al., 2019], it is distinct from the other GBM cell states. Interestingly, recent work has demonstrated that the MES-like state is driven by interactions between the tumor cells and immune cells, and has suggested that the targeted induction of the MES-like cell state together with immunotherapy may represent a novel opportunity for therapeutic intervention [Hara et al., 2021]. The neurodevelopmental-like transcriptional modules (NPC-/OPC-/AC-like) were more phylogenetically correlated with each other than any individual module was with the MES-like module (**Fig. 6E**). However, among the neurodevelopmental transcriptional modules, the AC-like module was the most phylogenetically correlated with the MES-like module, suggesting that transit between neurodevelopmental-like (NPC-/OPC-/AC-like) and MES-like states is driven by the AC-like state. To explore these relationships between GBM states further, we next used the phylogenetic correlations of GBM cell states, as determined by the per cell maximum transcriptional module scores, to infer cell state transition probabilities. This analysis revealed that stem-like cells primarily differentiated into AC-like cells, which could either dedifferentiate back into a stemlike state [Chaligne et al., 2021] or progress to the MES-like state (**Fig. 6F**). Notably, this inference suggests that, in this patient, the MES-like state derives from transitioning AC-like cells. This observation is consistent with recent findings that show that many MES-like cells have AC-like properties [Chanoch-Myers et al., 2022] and that the recep-tors (*e.g*., OSMR, EGFR, PDGFRB, and AXL) for ligands that drive transition into the MES-like state are expressed in AC-like cells but not stem-like cells [Hara et al., 2021]. PATH transition inferences from another human patient-derived GBM sample MGH122, from Chaligne et al. [2021], agreed with inferences from MGH115, revealing that of the neurodevelopmental-like cell states, AC-like cells appear to transition to the MES-like state at the highest rate (**Fig. S6A**).

To experimentally corroborate these cell state transition inferences obtained from primary human samples, we leveraged the artificial Molecular recorder approach [Chan et al., 2019] to trace gliomasphere phylogenies, using MGG23 [Wakimoto et al., 2011], a human patient-derived gliomasphere model (**Methods: Gliomasphere phylogenies**, **Fig. 6G**). Gliomaspheres are spheroid GBM cultures capable of recapitulating parental tumor cellular diversity [Laks et al., 2016], and thus represent an appropriate setting to measure cell state heritability versus plasticity. Two gliomasphere MGG23 replicates were grown *in vitro* for 4 weeks, at which point phylogenies were reconstructed using recovered barcodes, and cells were annotated according to their scRNAseq profiles. Consistent with the human patient data (**Fig. 6E**), PATH measurements in the gliomasphere model also showed higher phylogenetic correlations between the neurodevelopmental-like modules, than between any of the neurodevelopmental-like and MES-like modules (**Fig. 6G**). Furthermore, among the neurodevelopmental-like modules, the AC-like module was, as in patient sample MGH115, the most correlated with the MES-like module. Thus, using both native and artificial approaches for phylogenetic tracing in primary human samples and an *in vitro* model, respectively, we observed a strong phylogenetic relationship between the AC- and MES-like transcriptional programs; consistent with a model in which the MES-like cell state primarily derives from the AC-like state.

Finally, after analyzing the heritability of predefined glioblastoma gene transcriptional modules, using gene set enrichment analysis (GSEA) [Subramanian et al., 2005] we next profiled the heritability of the 3,000 most variably expressed genes in MGH115 (**Table S4**), ranked by their autocorrelation z scores, to discover heritable modules in an unbiased fashion. Consistent with **Fig. 6D**, this revealed an overrepresentation of five (NPC1/OPC/AC/MES1/MES2) GBM gene modules. This analysis further revealed that targets of the Polycomb repressive complex 2 (PRC2) constituents (*i.e*., targets of EED, SUZ12, EZH2), as well as sets of genes with promoters characterized by high CpG density and the repressive histone mark H3K27me3, in multiple stem cell contexts, were also enriched among heritably expressed genes in glioblastoma (**Fig. 6H, Table S5**). Similarly, brain tissue genes with bivalent promoters that are dually marked by both H3K27me3 and the activating mark H3K4me3, were also enriched among heritably expressed genes (**Fig. 6H**). This promoter methylation pattern represents a poised functional state that generally resolves to repressed (H3K27me3-only) or active (H3K4me3-only) states as cells differentiate. Promoter H3K27me3 levels are maintained primarily by targeting of the chromatin modifying PRC2, preventing differentiation by repressing lineage-specific gene expression [Boyer et al., 2006]. Notably, activity at PRC2-targeted sites is a key switch in the differentiation and maintenance of glioma stem cells [Natsume et al., 2013, Suvà et al., 2009].

To understand the relationships between these highly heritable gene modules, we next analyzed the enrichment of gene sets within distinct heritable gene modules defined by cross-correlations, with Over-Representation Analysis (ORA) [Korotkevich et al., 2021]. Hierarchical clustering of the phylogenetic correlations between the top 100 most auto-correlated genes revealed two heritable gene modules in MGH115 (**Fig. S6B**, **Table S6**). The first heritable module was enriched for gene sets associated with the neurodevelopmental-like glioma cell states (NPC1/OPC/AC), EED (a PRC2 subunit) target genes, and genes with high CpG density promoters with H3K27me3. This result is consistent with our previous observation that PRC2-target genes are preferentially hypomethylated, accessible and activated in the stem-like cell states [Chaligne et al., 2021]. The second heritable module was enriched for genes associated with the MES-like state and gene signatures associated with hypoxia. These results suggest that in patient MGH115, glioblastoma cells could occupy one of two heritable transcriptional states, either neurodevelopmental-like or mesenchymal-like. Cells could transit between these two states, primarily when occupying the more astrocyte-like end of the neurodevelopmental-like spectrum. Further, the neurodevelopmental-like module, in particular the stem-cell like states, is likely heritably maintained by PRC2 activity. These findings further highlight PATH’s ability to extract epigenetically grounded and biologically relevant expression profiles from single cell transcriptional and phylogenetic data in an unbiased manner.

In summary, the application of PATH to primary human glioblastoma samples identified the expected phylogenetic similarity by spatial location, nominated AC-like cells as the candidate precursor for MES-like cells, and highlighted the role of PRC2 in stable propagation of stem-like cell states. Thus, PATH can provide critical insight as to the biology underlying transcriptional cell state diversity in cancer.

### Quantifying cell state heterogeneity in B-cell acute lymphocytic leukemia (B-ALL) using single-cell whole genome sequencing

An exciting next frontier in the analysis of somatic evolution in humans is using somatic mutations as native lineage barcodes for lineage tree reconstruction from singlecell whole genome sequencing (scWGS). Current approaches often rely on costly and low-throughput single-cell cloning followed by WGS [Lee-Six et al., 2018], as somatic mutation rates are low and many scWGS methods suffer from high error and dropout rates, impacting the ability to call somatic variants with high confidence from single cells. To circumvent these challenges, and to explore PATH application to newly generated single-cell phylogenies constructed from the whole genome sequencing of single cells, we harnessed primary template-directed amplification [Gonzalez-Pena et al., 2021], a scWGS method based on a quasi-linear amplification that allows for high reproducibility and low allelic dropout. We aimed to construct a high-resolution lineage tree from scWGS of a B-ALL patient sample (**Fig. 7A**) with accompanying flow cytometry data for cell surface markers, and then apply PATH to determine the heritability versus plasticity of therapeutically relevant phenotypes in tumor cells.

**Figure 7:**
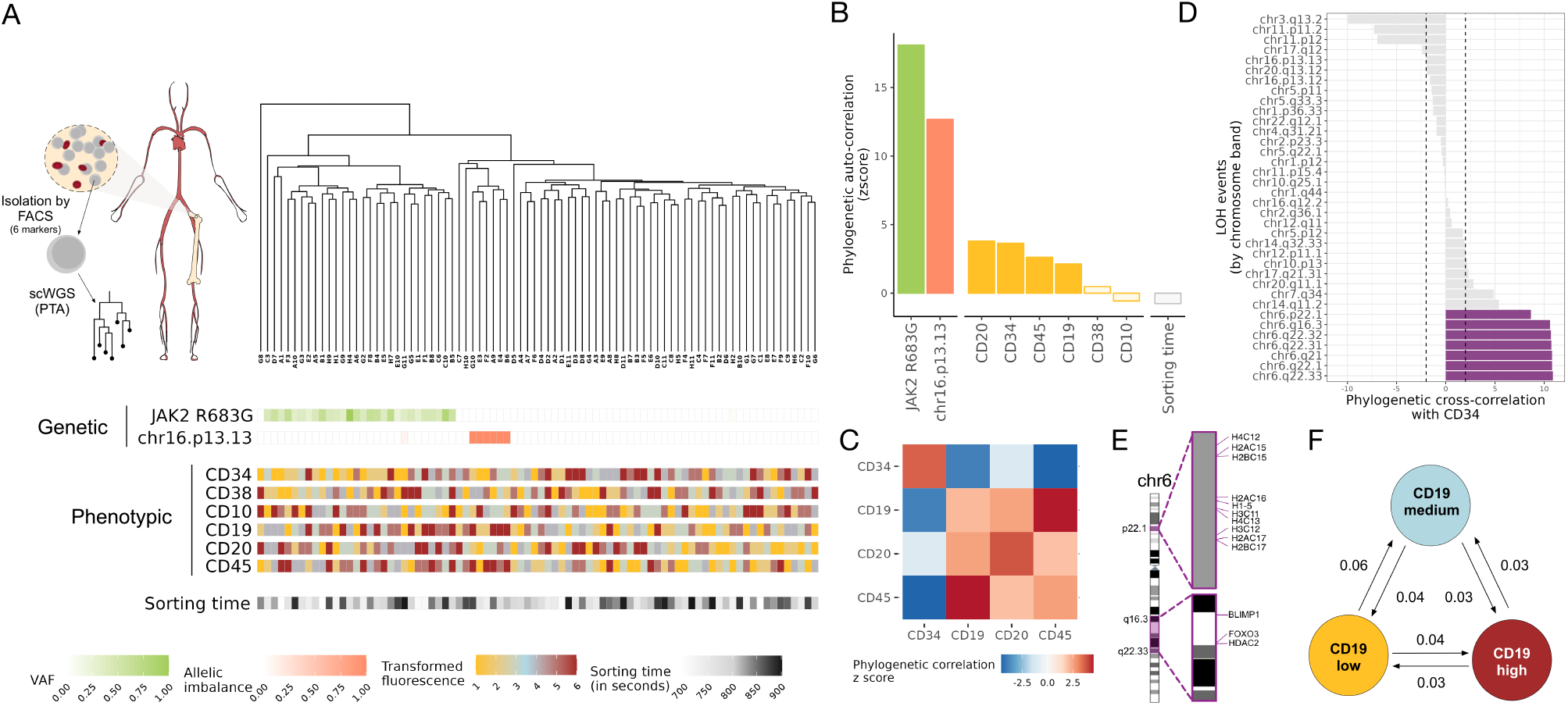
Quantifying cell state heterogeneity in B-ALL using single-cell whole genome sequencing. **A**) Top left-schematic of single-cell whole genome sequencing (scWGS) by primary template-directed amplification (PTA) of bone marrow isolated B cells sorted using six cell surface markers from a B-cell acute lymphocytic leukemia patient. Single-cell whole genome sequences were used to construct a single-cell phylogeny. Top right-Lineage tree constructed from single-cell whole genome sequences from a B-ALL patient sample (N=82 cells; ~8x coverage). Bottom-Genetic [allelic imbalance of germline heterozygous SNPs indicating a copy-number deletion at chr16; variant allele frequency (VAF) of single-nucleotide variant (SNV) of *JAK2*], Phenotypic (fluorescence of cell surface markers) and Random (sorting time) traits projected onto leaves. Cell surface markers used for cell sorting: CD34, CD10 and CD38 represent more immature B cell states, CD19, CD20 and CD45 represent more mature B-cell states. **B**) Phylogenetic auto-correlation z scores for genetic (copy-number deletion and SNV as in **B**), phenotypic (cell surface protein markers) and random (sorting time) factors. **C**) Phylogenetic correlation z score heat map for heritable cell surface protein markers. **D**) Phylogenetic cross-correlation z scores for CD34 and copy number deletions. Phylogeny annotated with genome-wide copy number deletion map can be found in **Fig. S7**. **E**) Chromosomal regions of deletions in clones with high CD34 expression. **F**) PATH inferred transition probabilities between states (CD19 low, medium and high) using all cells. Values rounded to the nearest hundredth.

To leverage somatic mutations as native lineage barcodes, we generated whole genome sequences for 86 cells (~8x coverage) sampled from a patient with B-ALL (**Methods: BALL analysis**) and quantified levels of cell surface markers that represent both more immature B cell states (CD34, CD10 and CD38) and more mature B cell states (CD19, CD20 and CD45) [Welner et al., 2008]. We used 55,251 single nucleotide variants (SNVs) to construct a high-resolution phylogeny (**Methods: B-ALL analysis**), annotated with genetic (copy number deletion, exonic SNVs excluded from tree reconstruction) and phenotypic (cell surface marker expression) information, with sorting time as a control for a random, non-heritable trait (**Fig. 7A**, **Table S7**). To determine the heritability of each trait, we applied PATH to these data to calculate phylogenetic correlations. As expected, genetic variation was highly heritable and sorting time, a random control, was not heritable (**Fig. 7B**). However, the phenotypic information was more variable; the majority of markers had intermediate phylogenetic scores that were between those of the genetic and random traits, with CD34 and CD20 displaying the highest heritability (**Fig. 7B**). These results showed that PATH can be used to analyze single-cell phylogenies generated from scWGS data and to measure the heritability of cell-surface protein expression markers in tumor cells.

To more deeply explore the biology of these tumor phenotypic traits, we next calculated the phylogenetic crosscorrelation between the significantly heritable cell surface markers (**Fig. 7C**). PATH showed that a marker associated with more immature B cells (CD34) negatively crosscorrelated with markers associated with more mature B cells (CD19, CD20 and CD45), which in turn were strongly crosscorrelated with one another. These results indicated that this B-ALL sample comprised tumor cells with heritable earlier and later B cell states, suggesting that some structure of the normal B cell differentiation trajectory is retained in this sample.

Taking advantage of the multimodality of the single-cell lineage data, we next sought to identify genetic features that correlated with CD34 expression, a marker that displayed high heritability and that reflects a more immature B cell state. To associate genetic and phenotypic features, we calculated phylogenetic correlations between copy number deletions and CD34 expression. PATH identified high phylogenetic correlations between CD34 expression and chromosome 6p22.1 and 6q16-q22 region deletions (**Fig. 7D, Fig. S7**), indicating that tumor clones that harbored these specific deletions also had higher CD34 expression. To identify potential genetic contributors that are associated with CD34 expression in these tumor clones, we more closely analyzed the deleted chromosomal regions and their impacted genes. Interestingly, these regions harbor genes that encode important B cell differentiation factors including PRDM1, FOXO3 and HDAC2 on 6q, as well as a histone gene cluster on 6p (**Fig. 7E**). Notably, it has been shown in B-cell lymphoma that deleterious mutations in histones H1B/H1-5 can cause remodeling of the chromatin state [Yusufova et al., 2021], leading to expression of stem cell genes, which is consistent with the earlier B cell state phenotype that we observed in cells harboring these deleted regions in this BALL sample. Therefore, it is possible that copy number loss of these regions and deletion of these genes could potentially contribute to the emergence of an earlier, more stem cell-like state (CD34 high). Indeed, 6p22.1 is known to be relatively frequently deleted in B-ALL and 6q16-q22 in DLBCL [Brady et al., 2022, Chapuy et al., 2018], further supporting the link between these deletions and a more stem-like state in this sample. Thus, PATH showed that quantifying the heritability of phenotypes and analyzing cross-correlation with genotypic features nominates candidate genotype-to-phenotype associations.

Finally, we sought to harness the ability of PATH to quantify transition dynamics between cell states to interrogate the plasticity of B-ALL targets of immunotherapy. In contrast to acute myeloid leukemia, where tumor cells develop from a more restricted window of cells from across the hematopoietic developmental trajectory [Miles et al., 2020, Zeng et al., 2022], B-ALL is considered more functionally plastic based on transplantation assays [Rehe et al., 2013] and cell-of-origin studies [Johnsen et al., 2014]. However, there is limited direct evidence of lineage-informed cell state plasticity and transitions directly in human samples at the single-cell level. Importantly, B-cell markers including CD19 have been used as targets for chimeric antigen receptor T (CAR-T) cell therapy [Davila and Brentjens, 2016, Maude et al., 2014], and while this approach has had success, there remain limitations in efficacy and sustained response [Schroeder et al., 2022]. B-ALL relapse after treatment with CD19-targeted CAR-T cells can be driven by genetic loss of CD19 [Xu et al., 2019], but other mechanisms, including the intrinsic plasticity of cell states associated with CAR-T target expression, could affect treatment implementation and success. We note that while PATH showed that CD19 expression had positive phylogenetic auto-correlation, (**Fig. 7B**), this marker had lower heritability compared to other analyzed markers and was substantially lower than the heritability of genetic traits, suggesting that CD19 expression was at least partially plastic. Indeed, PATH quantification of the transitions between high, medium and low CD19 expression states (**Methods: B-ALL analysis**) showed that while CD19 expression states were largely stable, we detected transitions between all three states. In particular, the low CD19 expression state was more likely to transition to the medium state, while the high CD19 expression state was about equally likely to transition to medium or low states (**Fig. 7F**). Thus, these results showed that there is a low level of fluid transitions between high, medium and low CD19 states, suggesting that in this B-ALL sample, while CD19 expression was a heritable trait with a positive phylogenetic correlation, it also exhibited a degree of plasticity between these expression level states. Altogether, these results and analyses highlighted the power of single-cell whole genome sequencing for phylogenetic analysis of human tumor cells, as well as the ability of PATH to quantify the heritability of therapy-relevant traits in a lineage-informed manner in order to gain insights into the plasticity of tumor cell states across subclones of a phylogeny.

## Discussion

The cells that comprise a multicellular organism derive from a single ancestral cell, thus remaining nearly genetically identical. Despite this genetic similarity, somatic cells within a multicellular organism encompass vast functional and phenotypic diversity. This phenotypic diversity can be maintained across mitotic divisions through the heritable transmission of both cell-intrinsic factors, such as epigenetic marks [Bintu et al., 2016, Halley-Stott and Gurdon, 2013] (*e.g*., DNA methylation and histone modifications) and cell-extrinsic factors (*e.g*., microenvironment). Each somatic cellular division, however, presents an opportunity to introduce changes to these heritable factors, for example in the form of heritable genetic or epigenetic changes. The phenotypic effect of these changes, however, is highly context dependent. In the case of cancer, mutations in putative cancer driver genes do not always lead to tumorigenesis and depend on cellular identity. For example, the malignant competence of BRAF mutations is dependent on the transcriptional background [Baggiolini et al., 2021], and some somatic mutations that confer a proliferative advantage are masked when found in progenitor cells [Nam et al., 2019]. As the presence of phenotypic variation provides a substrate for natural selection, an understanding of how these phenotypes are differentially encoded and inherited will help us dissect how cells in the soma evolve throughout the lifespan. To achieve this, however, we need an integrative model of somatic evolution informed by phenotypically annotated phylogenies. As such, scRNAseq is not sufficient and must be coupled with technologies that can also deliver information on cell ancestry.

To address this gap, PATH delivers an analytic framework needed for analyzing novel multi-omic lineage tracing single-cell datasets. PATH achieves this by building upon approaches from quantitative genetics and evolutionary biology used to measure heritability and phylogenetic signal [Blomberg and Garland, 2002] and adapts these to a somatic context. Specifically, PATH offers a bivariate generalization of phylogenetic signal in the form of phylogenetic correlation. Using phylogenetic correlations, PATH measures the ancestral dependency of single-cell phenotypes to infer their heritability versus plasticity. Additionally, for categorical phenotypes, such as a cell state or identity, PATH can transform phylogenetic correlations into state transition probabilities and thus allows for the inference of unobserved cellular dynamics. Importantly, this transformation also makes the classic interpretation of phylogenetic signal more concrete, as phenotypic transition dynamics are directly linked with the measurement of phylogenetic signal.

In step with the rapid advancement of lineage tracing technologies, other frameworks, such as *Hotspot* [Detomaso and Yosef, 2021] and *The Lorax* [Minkina et al., 2022], have been developed to study the lineage dependency of phenotypes in the single-cell context. Unlike other approaches, however, PATH can connect such measurements with a model of evolutionary dynamics and infer (categorical) phenotypic transition probabilities. Leveraging this connection, PATH allowed us to study how technical (*e.g*., sampling and reconstruction fidelity) and biological variables affect heritability measurements. This can inform our interpretations, for example, as PATH makes it clear that when sampling is sufficiently sparse, heritable phenotypes will likely appear plastic.

Other methods have also been advanced to estimate state transitions from phylogenies. For instance, if representing phenotypic (*e.g*., cell type) transitions as a Markov model, transition probabilities can be fit using Maximum Likelihood Estimation (MLE) [Louca and Pennell, 2019] or inferred with kin correlation analysis (KCA) [Hormoz et al., 2015, 2016]. PATH’s inference approach is more akin to KCA, as it transforms correlations into transitions; however, PATH can additionally be applied to subsampled phylogenies and when branch length measurements are absent. MLE, on the other hand, is commonly used in evolutionary biology to infer phenotypic transitions from species phylogenies. This approach takes the structure of the entire phylogeny into account (as opposed to just phylogenetic correlations) and searches for optimal transition rates. PATH’s accuracy is comparable to MLE, but computationally faster, particularly for larger trees with many phenotypes. This ability to accurately handle large trees with speed renders PATH suitable for analyzing single-cell phylogenies, which often contain many states, and an ever growing number of cells.

Using PATH, we studied previously published developmental lineage tracing datasets in early stages of embryological development [Chan et al., 2019] and brain organogenesis [Raj et al., 2018]. In murine development, we were able to analyze phylogenetic correlations between the blastocyst, the germ layers and specialized tissues, reconstructing known developmental trajectories and importantly, capturing the dual origin of the gut endoderm from both the epiblast and primitive endoderm [Kwon et al., 2008, Rothová et al., 2022, Saykali et al., 2019], which would not be achievable with scRNAseq alone. This highlights the ability of PATH to distinguish between phenotypic and ancestral similarity. We further showed that, consistent with a model of epigenetic inheritance and our understanding of imprinting throughout development [Loda et al., 2022], a unique X-chromosome expression profile is inherited by gut cells with extraembryonic origins. In zebrafish brain development, we used PATH to show how anatomic proximity influences relatedness of neurons in the developing brain and further highlighted PATH’s ability to coordinate transcriptional and anatomic data to show a shared lineage between substructures in the fore, mid and hind brain. As multi-modal singlecell technologies improve, PATH could be applied to coordinate transcriptional data with other modalities, beyond anatomic location, to interrogate fundamental questions in development. We also observed a striking pattern of stable lineage commitment for both excitatory (glutamatergic) and inhibitory (GABAergic) neurons in the forebrain. As lineage tracing techniques improve, using PATH we may eventually be able to more finely map the transitions undergirding cell state differentiation hierarchies in these functionally complex organs and reveal the factors responsible for maintaining and modifying lineage commitments.

Many scRNAseq analyses have revealed cell state diversity in cancer, but representing only a snapshot, have been unable to determine how temporally stable or transient such cell states are. Using PATH on lineage traced scRNAseq data, we can bypass this constraint, to quantify cell state temporal dynamics. To demonstrate this potential, we applied PATH to two previously published single-cell cancer datasets [Chaligne et al., 2021, Simeonov et al., 2021]. First, we observed that spatial location was highly stable: metastatic tissue location in a mouse model of pancreatic cancer, and tumor region in a human glioblastoma. Second, we used PATH to study transcriptional stability. It is not yet clear whether cancer cell state diversity predominantly reflects transient transcriptional fluctuations akin to entering and exiting the cell cycle, or more stable transcriptional changes analogous to cell fate commitment in development. In both cancer datasets, we observed the heritability of transcriptionally defined cell states in two of the largest drivers of cancer cell state diversity – position along the EMT continuum in pancreatic cancer, and in the stem cell hierarchy in glioblastoma. Interestingly, in both of these cancers, cell states were not uniformly plastic/heritable. Future application of PATH to other cancers could guide future treatments, such as the strategic targeting of specific transcriptional states, or the therapeutic modulation of state transition rates, in order to drive tumors to extinction.

Underscoring this potential, our analysis of newly generated data from a B-ALL patient demonstrated that using a powerful new single-cell whole genome sequencing approach (PTA) enabled construction of a high-resolution tumor cell phylogeny, and that application of PATH to this annotated tree yielded a detailed cancer profile encompassing genetic, phenotypic and ancestral dimensions. This PATH profile provided quantitative measurements of the heritability and plasticity of cell surface marker expression, revealing heritability of early vs. late B cell differentiation states, and linking these state biases with potential underlying genetic aberrations. Moreover, PATH analyses also quantified the plasticity of the therapeutically-relevant B-ALL marker CD19, which has been successfully used as a target of CART immunotherapy [Schroeder et al., 2022]. As cell state plasticity in the expression level of a therapeutic target can serve as a potential evolutionary therapeutic escape mechanism, we propose that such information could potentially serve to prioritize therapeutic targets for clinical development.

We speculate that as sequencing costs continue to fall, clinical single-cell whole genome sequencing for phylogeny reconstruction and analysis of tumor samples could become more accessible, rendering such approaches feasible.

In conclusion, somatic evolution represents an exciting frontier in evolutionary biology, where asexually reproducing somatic cells evolve over the multicellular organism’s life span. Studying this frontier requires analytical advances in step with technological advances that provide multi-modal single-cell annotation with high resolution phylogenetic information. We envision that PATH can thus help transform qualitative key concepts in multicellular somatic biology such as fate-commitment, heritability and plasticity into precise measurements, with broad impact on our understanding of organismal health and disease. As future technology evolves to capture phylogenetic information with epigenetic and spatial information, we further envision that the adaptability of the PATH framework will enable the linkage of cell state heritability and the mode of inheritance propagation (*e.g*., genetic, epigenetic, cell-extrinsic) to define the fundamental principles of somatic evolution.

### Limitations

Mathematical models represent an idealized situation, and in practice, can be robust to small violations to their assumptions. As outlined in the results and methods sections, several assumptions are made in PATH’s cell state transition inference model (*e.g*., transitions are Markovian, cell states are near their equilibrium proportions). These assumptions should be (nearly) met if transition rates only depend on a cell’s current and not prior states, and when sampling is not biased. Other assumptions, such that cell birth or death rates do not differ as a function of cell state, could be violated and impact inferences. Specifically, if some cell states have much higher proliferation rates than others, inferred transition rates could be biased. Such a scenario represents an opportunity for future model development. However, such a model would likely rely on accurate branch length measurements and higher resolution singlecell phylogenies than are typically available now. Transition inference accuracy is also most reliable when heritability is significantly detected, as demonstrated in **Fig. 2C,D**, and inferences from phylogenies with insignificant phylogenetic correlations should be interpreted cautiously.

Additionally, the robustness of PATH measurements is dependent on the quality and resolution of the lineage data, and analysis of sparsely sampled trees can lead to underestimation of heritability, as shown by our simulations. Relatedly, PATH is subject to the standard problems affecting single-cell analyses, including data dropout, accuracy of cell state assignment algorithms, completeness of gene set modules and batch effects. These limitations may constrain the analysis of currently available datasets; however, we anticipate that with advances in lineage tracing and singlecell multiomics technologies, PATH’s utility will expand as single-cell lineage tree data continue to improve.

Most single-cell phylogenies do not include branch length estimates, which can further confound inferences. PATH, however, was designed to accommodate some of these limitations, by imputing branch lengths, and by focusing on closer (one-node apart) phylogenetic relationships.

As more multi-omic single-cell lineage tracing experiments are conducted, and lineage tracing and other technologies further mature, allowing for even higher resolutions of phylogenetic relationships and phenotypic states, more subtle evolutionary dynamics could be teased apart with PATH. If multiple layers of information, in addition to transcriptional phenotype and ancestry, such as location or microenvironment, are gathered for each cell, measured phylogenetic correlations across these layers could help dissect the encoding of heritable phenotypes. That is, phylogenetic correlations between phenotypes and microenvironments could help determine whether a heritable phenotype is encoded intrinsically (*e.g*., via genetic or epigenetic mechanisms) or extrinsically (*e.g*., via shared microenvironment stimuli).

### Conclusion

In summary, throughout a multicellular organism’s lifetime, its constituent somatic cells continuously evolve, accumulating heritable phenotypic variation. When positively selected, heritable phenotypic variation deleterious to the organism as a whole may also lead to disease states or malignancy, which itself represents a “runaway” evolutionary process. PATH formally connects the analysis of cell state diversity and somatic evolution, and quantifies critical aspects, replacing *qualitative* conceptions of “plasticity” with *quantitative* measures of cell state transition and heritability. The application of PATH thus powerfully brings together approaches from evolutionary biology and single-cell technology, to study complex dynamics governing somatic evolution – an exciting novel frontier in multicellular biology.

## Acknowledgments

We thank members of the Landau laboratory and Norbert Fehér for thoughtful discussions throughout the development of this work. We thank Nir Yosef for critical comments on the manuscript. We thank Aaron McKenna and Bushra Raj for sharing data and code related to the scGESTALT phylogenies. We thank Alexander Meissner’s group and the authors of Chan *et al*. 2019 for sharing their cell type assignment data and code. CG is supported by a Burroughs Wellcome Fund Career Award for Medical Scientists, National Institutes of Health Director’s New Innovator Award (DP2-CA239145), and Chan Zuckerberg Investigator Award. DAL is supported by the Burroughs Wellcome Fund Career Award for Medical Scientists, the Valle Scholar Award, the William Rhodes and Louise Tilzer-Rhodes Center for Glioblastoma at NewYork-Presbyterian Hospital (NYPH 203205-01), the Sontag Foundation (Distinguished Scientist Award, SFI203261-01), the National Institutes of Health Director’s New Innovator Award (DP2-CA239065), Leukemia Lymphoma Scholar Award and the Mark Foundation Emerging Leader Award. This work was supported by the National Heart Lung and Blood Institute (R01HL157387-01A1), National Cancer Institute (R01 CA242020, R01 CA251138, and P50 254838), a Tri-Institutional Stem Cell Initiative award and the National Human Genome Research Institute, Center of Excellence in Genomic Science (RM1HG011014). DAL and MLS are jointly supported by NCI R01CA258763 and a grant from the STARR Cancer Consortium.

## Author contributions

JSS, ARD, TP and DAL conceived the project and designed the study. JSS developed PATH and performed simulations. ARD, JSS, TP and SR performed analyses. YF and TH generated the gliomasphere data. YP and CG generated the single-cell PTA data. JSS, ARD, TP, MLS, CG and DAL helped interpret the results. MLS, YF, CG and TH provided critical comments on the manuscript. JSS, ARD, TP, CP and DAL wrote the manuscript. All authors reviewed and approved the manuscript.

## Code availability

The code used to measure phylogenetic correlations and to infer cell state transitions is available as part of our *PATH* R software package at https://github.com/landau-lab/PATH. Code used for data processing and analysis will be made available upon publication.

## Methods

### Phylogenetic correlations

To quantify the distribution of a single-cell measurement, such as transcriptional state, across a phylogeny, we use Moran’s *I* [Moran, 1950], a classic measure of spatial auto-correlation. We also import its bivariate generalization, a measure of spatial cross-correlation [Chen, 2015, Wartenberg, 1985] to quantify pairwise phylogenetic cross-correlations [Chaligne et al., 2021]. For this study, we refer to both phylogenetic auto- and cross-correlations as *phylogenetic correlations*.

To compute the phylogenetic auto-correlation of a single variable (Moran’s *I*), we need a measurement of pairwise distances between cells, provided by the phylogeny, and a standardized observation per cell (with mean subtracted and normalized by population standard deviation).

For example, the expression of a particular gene in *N* cells could be represented by the *N*-dimensional vector *x*, where each element represents an expression score per cell. This vector is then standardized, producing the vector *z_x_* = (*x* – *μ_x_*) /*σ_x_*, where *μ_x_* and *σ_x_* are the mean and population standard deviation of *x*, respectively.

Pairwise phylogenetic distances (*e.g*., node or branch length distances), represented by the elements of the square *N*-dimensional matrix *L*, are transformed into a phylogenetic weight matrix *W*, with a chosen weighting function *f_w_*, such that *W* = *f_w_* (*L*). This function first weights each off-diagonal element of *L*, and then sets diagonal elements of *L* to 0. An example of a weighting function is the inverse of phylogenetic distance (*i.e*., for *i* ≠ *j*, *W_ij_* = 1/*L_ij_*, otherwise *W_ij_* = 0). Another example of a weighting function that we use throughout this study is to select only a specific phylogenetic distance (*e.g*., for *L_ij_* = *d* and *i* ≠ *j*, *W_ij_* = *L_ij_*, otherwise *W_ij_* = 0), where *d* is either a chosen branch or node distance. These weights are then normalized such that they sum to 1, resulting in a normalized weight matrix, 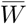. The phylogenetic auto-correlation of *x* is then defined as,

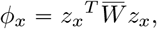

where superscript *T* signifies the matrix transpose.

The phylogenetic cross-correlation between two different single-cell measurements (bivariate Moran’s *I*), is calculated similarly, where both *z_x_* and *z_y_* are standardized single-cell measurements or observations corresponding to the vectors *x* and *y*,

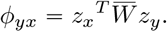

All pairwise phylogenetic (auto- and cross-) correlations can be computed simultaneously if single-cell measurements are in matrix form. Single-cell measurements are represented by the *N* × *n* dimensional matrix *X*, in which its *N* rows represent individual cells and its *n* columns represent distinct measurements (such as the expression of *n* distinct genes). When measuring phylogenetic correlations for a categorical states, in which a cell can occupy only one of a set of possible states at any given time (*e.g*., cell type), each column of *X* denotes a distinct cell state, and the state of each cell is indicated by a 1 in the appropriate column, and 0s in the remaining columns. For example, if the *i*th cell is in the second of two possible cell states, then *X*_*i*,1_ = 0, and *X*_*i*,2_ = 1. For all measurement types, the columns of the single-cell measurement matrix *X* are standardized, as above, to produce the *N* × *n* dimensional matrix *Z*, which is then used to compute the square *n*-dimensional phylogenetic correlation matrix,

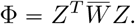

Note that the diagonal elements of Φ correspond to phylogenetic auto-correlations. Furthermore, phylogenetic correlation z scores can be calculated by performing a leaf-permutation test or analytically with moments from Czaplewski and Reich [1993]. Phylogenetic correlations and analytical z scores can be computed with the function `xcor()` in our `R` software package. Additionally, normalized phylogenetic weight matrices can be computed using either `one_node.tree.dist()`, `inv.tree.dist()`, or `exp.tree.dist()` from our *PATH* R package.

Note that phylogenetic correlations depend on the structure of the matrix 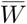, thus weighting functions should be chosen carefully. For the purposes of this study, we predominantly use a weighting function that only includes cells that are each other’s nearest phylogenetic neighbor, specifically cells that are separated by a node distance of one.

### Simulating phylogenies

In this study we use two approaches to simulate single-cell phylogenies. We simulate *idealized phylogenies*, which are completely sampled, discrete-time, bifurcating, ultrametric, and balanced phylogenies that contain *N* = 2^*g*^ cells, where *g* is the number of generations that have occurred since the root. Additionally, each branch length, which corresponds to one generation, has a length of one. To generate an idealized phylogeny we use the function `pbtree`(`b = 1, d = 0, n = N, type = “discrete”`) from the R software package *phytools* [Revell, 2012].

We also simulate phylogenies using what we refer to as a *sampled somatic evolutionary process*, which is a sampled and continuous-time birth–death process, using the function `generate_tree_hbd_reverse()` from the R software package *castor* [Louca, 2020, Louca and Doebeli, 2018]. In contrast to idealized phylogenies, these phylogenies can be imbalanced, and contain any number of cells that represent a fraction of the total somatic population. For these simulations, parameters for cell division (or birth), and cell death, the sampling rate, and the total number of sampled cells can be specified. Here, phylogenetic branch lengths correspond to time in continuous units, and not to generations, as in idealized phylogenies.

Cell state transition dynamics are represented as a discrete-or continuous-time Markov model (**Methods: Markov model of cell state transitions**) on idealized, and sampled somatic evolutionary phylogenies, respectively. Markov cell state transitions are simulated on both types of phylogenies using the *castor* function, `simulate_mk_model()`.

### Markov model of cell state transitions

We model cell state transition dynamics as a Markov chain [Grimmett and Stirzaker, 2020], in both discrete- and continuous-time.

For a discrete-time Markov chain comprising *n* possible cell states, the transition probabilities (corresponding to one unit of time) are stored in a square *n*-dimensional *transition matrix*, *P*. Individual elements of the transition matrix are referred to by their subscript coordinates, such that *P_ij_* refers to the transition probability located in row *i* and column *j* and represents the probability of switching from state *i* to state *j*. The probability that a cell in state *i* transitions to state *j* after *t* discrete time-steps is given by 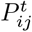 (note: superscript *t* reflects matrix, not element-wise, powers). As elements represent probabilities, each row of *P* must sum to 1.

Discrete-time chains might be more intuitive when recording times in non-overlapping generations, and continuous-time might be more appropriate when generation times vary and/or overlap. A continuous-time Markov chain has a *transition rate matrix*, *Q*. Each element, *Q_ij_* records the infinitesimal transition rate between states indexed by their row and column. The transition probability matrix can be recovered by matrix exponentiating the rate matrix, that is *P* = exp(*Q*), and the transition probability of switching from state *i* to state *j* after a (continuous) *t* amount of time is given by *P*(*t*) = exp(*Qt*). Lastly, each row of *Q* must sum to 0.

The *stationary distribution* of a Markov chain, if also a *limiting distribution*, represents the expected frequencies of each cell state at equilibrium, and is represented by the *n*-dimensional vector *π*. For large *t*, the transition matrix *P^t^*, if it has a limiting distribution, converge to the matrix Π, where each row of Π is equivalent to the vector *π*. This means that after a sufficiently long amount of time, the probability of transitioning from any state to state *j* is equal to state *j*’s equilibrium frequency, *π_j_*. For chains with symmetric transitions, where transitions to and from a state occur with equal probability (*i.e*., *P_ij_* = *P_ji_*), the equilibrium frequency for each state is 1/*n*, where, recall *n* is the number of possible cell states.

Finally, Markov chains are *reversible* if the products of the transition probabilities between two states and their stationary frequencies of origin are the same, *i.e*. *π_i_P_ij_* = *π_j_P_ji_*. Note that the reversibility of a Markov chain does not imply that transitions are symmetric, and that asymmetric Markov chains can also be reversible.

We connect Markov cell state transition dynamics with phylogenetic correlations in **Phylogenetic correlations and cell state transitions**, and use this connection to infer cell state transition dynamics from phylogenetic correlations in **Inferring cell state transitions from phylogenetic correlations**.

### Phylogenetic correlations and cell state transitions

Phylogenetic auto-correlations measure the phenotypic similarity of closely versus randomly related cells (with respect to ancestry). More generally, the phylogenetic crosscorrelation of two phenotypes, is a measure of the relationship between those phenotypes in closely related, as compared to, randomly chosen cells (**Methods: Phylogenetic correlations**). When measuring categorical states on phylogenies, if we use a phylogenetic weighting function that retains only specified phylogenetic distances and omits all others, phylogenetic correlations measure the difference between *state-pair* frequencies in closely (as specified by the retained distances) versus randomly related cell pairs. Here, *state-pair* refers to the states represented in a pair of chosen cells.

For example, on idealized phylogenies (**Methods: Simulating phylogenies**), if we apply a phylogenetic weighting function that preserves all branch lengths equal to two, and sets all other phylogenetic distances to zero, the phylogenetic correlation between two states will be a measure of the difference between the frequencies at which pairs of states are found within sisters versus random cell pairs. On idealized phylogenies, sister cells are separated by a branch length of two, because the branches that connect each of them to their parent, represent one generation, and thus have a branch length of one. Similarly, if a weighting function that retained only branch lengths equal to four is used, the resultant phylogenetic correlations, for an idealized phylogeny, would measure the difference between state-pair frequencies in first-cousins versus random cell pairs. In general, if we use a weighting function on an idealized phylogeny that only retains phylogenetic branch lengths equal to 2*t*, phylogenetic correlations would measure the difference between the frequencies at which specific state-pairs are found within pairs of cells that share a most recent common ancestor (MRCA) *t* generations ago, versus randomly chosen cell pairs (with replacement).

To illustrate, consider an idealized *N*-cell phylogeny and *n* possible cell states, in which the pairwise phylogenetic branch lengths between cells, represented by the square *N*-dimensional matrix *L*, and each cell’s categorical state, recorded in the *N* × *n* dimensional matrix *X* (as in **Methods: Phylogenetic correlations**), are known. First, a weighting function that only retains phylogenetic branch lengths equal to 2*t* is applied, such that *W*(*t*) = *f_w_* (*L,t*), and the sum of the weights in *W*(*t*) are normalized to equate to 1, resulting in the normalized phylogenetic weight matrix 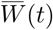. The frequency in which cells phylogenetically separated by a branch length distance of 2*t* are in states *i* and *j* is given by the *ij*th element of the square *n*-dimensional frequency matrix,

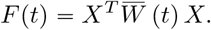

Note, that on a phylogeny, because the order of the cells within a pair is arbitrary, for *i* ≠ *j*, the frequency of observing either the state-pair *ij* or state-pair *ji*, is given by the sum of the frequencies *F* (*t*)_*ij*_ + *F*(*t*)_*ji*_. Additionally note that in the specific context of idealized phylogenies, state-pair frequencies as in *F*(*t*) are equivalent to *kin correlations* [Hormoz et al., 2016].

These state-pair frequencies can be transformed into phylogenetic correlations, Φ(*t*), by first subtracting the random (with replacement) state-pair frequencies, and then normalizing by the cell state population covariances, where *μ* and *σ* are the respective *n*-dimensional state frequency and population standard deviation vectors (and division is elementwise),

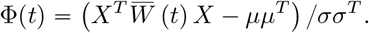

If cell state does not depend on ancestry, then we would not expect state-pair frequencies to substantially differ in closely and randomly related cells, resulting in low (near zero) phylogenetic correlations. However, if cell states can be inherited, but also sometimes stochastically transition, we would expect phylogenetic correlations to be generally non-zero. This is due to the fact that, if heritable, the states for cells that share a MRCA *t* generations ago will each depend on the state of the same ancestral cell. As such, state-pair frequencies and therefore phylogenetic correlations as measured above, will depend on how heritable each cell state is, and how often each state transition to another state occurs. In other words, the difference between state-pair frequencies in closely related versus random cells, might be attributable to underlying cell state transition and inheritance dynamics. To make this more concrete, below we link a Markov model of cell state transition dynamics with cell state phylogenetic correlations.

For cell state transition dynamics that can be represented as a Markov chain (**Methods: Markov model of cell state transitions**), we can predict state-pair frequencies for a given pairwise phylogenetic distance, from the transition probabilities *P* (a square *n*-dimensional matrix, where *n* is the number of cell states) and the limiting distribution *π* (an *n*-dimensional vector). For an intuitive example, consider the situation where a pair of sister cells (that share a parent) are in the same specific state. One way sister cells can end up in the same state is by both inheriting the same parental state, and subsequently not transitioning to another cell state. Alternatively, if the sister cells did not inherit their current state, they could have each independently transitioned from the parent’s state to the same new state. The probability of observing sister cells in the same specific state is then determined by summing the probabilities for each different scenario that could lead to such an outcome. The probability of each scenario is computed by taking the probability that the unobserved ancestral cell (here the parent) was in a particular state, given by *π*, and multiplying by the relevant transition probabilities, provided by *P*. For the situation in which there are only two possible cell states, the probability of observing the state-pair *ij* (where one cell is in state *i* and its sister is in state *j*) is,

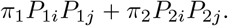

More generally, for *n* possible cell states, the probability of observing each state-pair (where one cell is in state *i* and the other is in state *j*, and *i* and *j* can range from 1 through *n*), in two cells that share a MRCA t generations ago, where *D* = diag(*π*) and superscript *T* is the matrix transpose, is

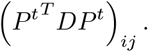

If the cell state transitions are reversible, then *P^T^D* = (*DP*)^*T*^ = *DP*, and the probability of observing each state-pair in cells separated by a phylogenetic distance of 2*t* can be simplified to be,

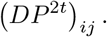

These equations show that, for Markov transition dynamics at equilibrium, the probabilities of observing each possible state-pair are determined by the probability that the shared ancestor was in a particular state, multiplied by the probability that such a state transitioned to the two descendant cell states observed *t* generations later, and then summed for each possible ancestral state. For reversible chains, this is also equivalent to the probability of starting in one of the descendant states, followed by a transition to the other descendant state after the 2*t* time-steps that separates them.

Using these equations, we can compute expected phylogenetic correlations for cell state transitions. This is achieved by subtracting the probability of observing randomly chosen cells (with replacement) from the state-pair probabilities and normalizing by the cell state covariances,

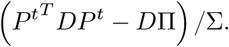

For reversible transitions, this simplifies to,

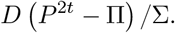

An illustration for these calculations for two cell states is depicted in **Box S1**. Notice that as *t* increases, *P*^2*t*^ → Π, and all phylogenetic correlations thus approach 0. This means that as cell pairs become more distantly related, their state-pair frequencies should approach those as if the two cells comprising the pair were drawn at random from the population. Also note that the closer transition probabilities are to cell state equilibrium frequencies, the less heritable cell states will appear. Furthermore, in this context, a high cell state phylogenetic auto-correlation would imply that the probability of transitioning to any other state is relatively low, and thus that the cell state is highly heritable.

In the context of species evolution, the auto-correlative method of measuring phylogenetic signal was not based on an evolutionary model, in contrast to signal metrics like Pagel’s *λ*, and thus considered more difficult to interpret biologically [Münkemüller et al., 2012]. Here, not only do we define a bivariate measure phylogenetic signal using phylogenetic correlations, but we illuminate a connection between the measurement of phylogenetic auto- and crosscorrelations with a model of evolutionary dynamics. This relationship with (categorical) phenotypic transitions thus clarifies the interpretation of what phylogenetic correlations measure. Finally, although we only make the connection explicit for categorical phenotypic states, phenotypic “covariance structures” (which will affect phylogenetic correlations) can be linked with a variety of evolutionary processes, including models for the evolution of continuous phenotypic states [Hansen and Martins, 1996].

The relationship between phylogenetic correlations and reversible cell state transition dynamics, can be used to infer unknown transition probabilities from phylogenetic correlations, as demonstrated in **Inferring cell state transitions from phylogenetic correlations**.

### Inferring cell state transitions from phylogenetic correlations

#### Idealized phylogenies

For reversible Markov chains with a limiting distribution (**Methods: Markov model of cell state transitions**) operating on idealized phylogenies (**Methods: Simulating phylogenies**, and **Phylogenetic correlations and cell state transitions**), transition probabilities can be inferred by converting phylogenetic correlations back into state-pair frequencies (not centered or normalized) and then dividing each row *i* by 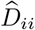, the corresponding cell state frequencies at a branch length distance of 2*t* (where 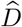 is an estimate of *D*),

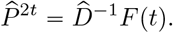

To arrive at the transition probabilities for a specific length of time, appropriate matrix powers or roots can be taken. For instance,

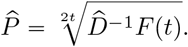

In this setting, using idealized phylogenies, this formulation is equivalent to inferring transition probabilities using *kin correlation analysis* (KCA) [Hormoz et al., 2016], and conceptually similar to an approach for approximating nucleotide substitution rates [Yang and Kumar, 1996].

Finally, note that in this context, if the Markov chain does not have a limiting distribution, for instance, if it is periodic, we might not be able to infer the correct transition probabilities. For example, in the situation where there are two possible cell states, and the transition probabilities to and from each state are *P*_12_ = *P*_21_ = 1, and the self-transition probabilities are *P*_11_ = *P*_22_ = 0, then the states of every observed cell (in the terminal generation) will be the same, but different from the states in the cells from the previous generation. For this case, we would correctly infer that the self-transition probability of the state observed in the terminal generation is 1 after 2*t* time-steps, however, our estimates for an odd number of time-steps would be incorrect.

#### Phylogenies from a sampled somatic evolutionary process

Phylogenies resulting from a sampled somatic evolutionary process (**Methods: Simulating phylogenies**) contain only a sampling of the somatic population under study and continuous and non-uniform branch lengths. These factors must be taken into account in order to successfully infer transition probabilities. To accomplish this, we take the state-pair frequency matrix (used to compute phylogenetic correlations) at a *node-depth* of *d*, *F*(*d*), by applying a weighting function that omits all phylogenetic distances that do not correspond to a node-depth equal to *d*, and the mean of the corresponding branch length distances *τ*. For each node-depth, we can approximate the transition matrix as,

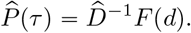

This is an estimate of the transition probability matrix for a time proportional to the mean branch length distance between cells *d* nodes apart. For a completely sampled idealized phylogeny, *τ* = 2.

More generally, we estimate *P*(*t*) (for time *t*), to be

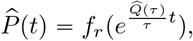

where 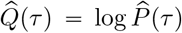, and *f_r_* () normalizes rows so that each sums to 1.

For circumstances in which branch lengths are unknown or inaccurate, for a node-depth of one, *τ* can be imputed if the cell sampling can be approximated and a model of somatic evolution is assumed. This can be accomplished by us-ing branch lengths from simulated phylogenies from our somatic evolutionary process (**Methods: Simulating phylogenies**), or approximated analytically (**Methods: Imputing branch lengths**). Cell state transition dynamics can be inferred with the function `PATH.inference()` in our R software package.

All inferred transition rates for the analyzed datasets were determined in this manner, using either 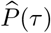 (as in **Figs. 6F, S6A**) or 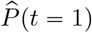 (as in **Figs. 4D, 5G, 7F**).

### Phylogenetic reconstruction

To simulate evolution, phylogenetic reconstruction, analysis and inference, we first simulate trees as a sampled somatic evolutionary process, a continuous birth–death process, (**Methods: Simulating phylogenies**) under various parameter schemes, in which the sampled tree size, and the birth, death, and sampling rates can vary. Once phylogenies are simulated, two distinct Markov processes are run: (1) a process simulating cell state transition dynamics, and (2) a process simulating the mutation/scarring of heritable cellular barcodes. The first Markov model is as described in the section **Markov model of cell state transitions**, and the second Markov model simulates barcode scarring and is a simple two-state, continuous-time, and symmetric model, with one rate parameter *s*, that runs independently for each mutable site contained within a cell’s heritable barcode.

The elements of the 2-dimensional square barcode scarring transition rate matrix are given by *Q*_11_ = *Q*_22_ = –*s*, and *Q*_12_ = *Q*_21_ = *s*.

Once both cell state transition dynamics and barcode mutations are simulated, a phylogeny is reconstructed – ignoring the true simulated phylogeny – with the unweighted pair group method with arithmetic mean (UPGMA) algorithm on pairwise-barcode Hamming distances. Branch lengths (evolutionary distances) are estimated from the number of barcode differences, using −0.5 log(1 – 2(*h*/*l*))/*s*, where *h* is the Hamming distance, *l* is barcode length, and *s* is the barcode cut rate.

Reconstructed phylogeny error is scored by computing the normalized Robinson-Foulds distance [Robinson and Foulds, 1981] and Mean Path Length distances [Steel and Penny, 1993] between the reconstructed and true trees. Phylogenetic correlations (using a node-depth of one weighting function) computed for the true and reconstructed tree are also compared by taking their mean differences. Lastly, transition inference is performed using two approaches (**Methods: Inferring cell state transitions from phylogenetic correlations**), by either using measured (determined by the Hamming distances) or imputed (**Methods: Imputing branch lengths**; determined using estimated parameters of a sampled somatic evolutionary process) branch lengths to derive 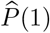 from 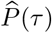. Accuracy for both methods is assessed by measuring the Euclidean distances between the inferred and true/simulated transition probabilities.

### Imputing branch lengths

For phylogenies in which branch lengths are unknown or potentially inaccurate, we can impute the phylogenetic branch lengths used to infer transition rates (**Methods: Inferring cell state transitions from phylogenetic correlations**) by using the sampled somatic evolutionary process model (**Methods: Simulating phylogenies**), using two approaches. In both cases, branch lengths are imputed by using either measurements or estimates to parameterize our sampled somatic evolution model. For the first, more exact, approach, we directly measure branch lengths that correspond to a node depth of one in simulations that use the estimated parameters. For the second, more approximate approach, we use an analytical expression, given a somatic evolutionary model parameterization, for computing the expected lengths of phylogenetic *pendant edges*, which are proportional to the branch length distances that separate cells phylogenetically one node apart. For a sampled somatic evolutionary process, pendant edge lengths are expected to be [Stadler and Steel, 2012],

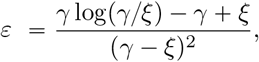

where *ξ* is the product of the cell birth and sampling rates, and *γ* is the net growth rate, given by the cell birth minus cell death rates. Using this expression, we can impute the approximate branch length distance between cells separated by one node, to be 2_*ε*_. For *γ* = 1 (where *ξ* is equal to the sampling rate, *N*_sample_/*N*_population_), as sampling becomes sparse, *ε* ≈ log (*N_population_*/*N_sample_*) – 1, and branch length distances at a node-depth of 1 are expected to be proportional the logarithm of the (inverse) sampling rate.

To test the robustness of our cell state transition inference approach when using imputed branch lengths, we input a sampling rate estimate by randomly selecting a rate within one order of magnitude above or below the true simulated rate. That is, if the simulated sampling rate was 10^-6^, we randomly select a sampling rate estimate between 10^-5^ and 10^-7^, for imputing branch lengths when inferring transition rates using PATH.

### Assessing cell state transition inference accuracy

To assess the accuracy of our inferences using PATH, we simulated phylogenies across a range of parameters, varying the cell sampling, birth and transition rates, as well as the number of cells and possible cell states. To generate a random *n*-dimensional transition rate matrix, for each cell state, (*n* – 1) numbers are drawn from a uniform random distribution, ranging between 0 and 0.1, and sequentially assigned to each off-diagonal matrix element per row. As rows must sum to 0, the remaining (diagonal) element in each row is set to the negative sum of these randomly drawn values. After parameters are chosen and a transition rate matrix is randomly generated, phylogenies are simulated (**Methods: Simulating phylogenies**) and phylogenetic correlations (**Methods: Phylogenetic correlations**) and inferences (**Methods: Inferring cell state transitions from phylogenetic correlations**) are computed.

We also compared cell state transition rate inference accuracy with MLE. To do this, we used the function `fit_mk()` from the R *castor* package [Louca, 2020, Louca and Doebeli, 2018] to estimate the transition rate matrix 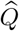 from a simulated phylogeny (**Methods: Simulating phylogenies**). To assess the accuracy of inferences using either PATH or MLE, we compute the Euclidean distance between the inferred transition probability matrix 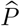, for *t* = 1, and the true transition probability matrix *P*. Inferences using both PATH and MLE were performed on the same simulated phylogenies, and accuracies compared.

### Mouse embryogenesis

Normalized RNA matrices and phylogenies were downloaded from Gene Expression Omnibus (GEO) series GSE117542 and imported into R (v. 4.1.3). Cell type annotations were provided upon request by the corresponding authors of the original publication [Chan et al., 2019]. Blastocyst layer annotations were inferred from germ layer membership. Phylogenies were extended by connecting node identifiers with single-cell barcodes using a dictionary provided in pickle files. We analyzed phylogenies for embryos 2 and 6 from [Chan et al., 2019]. Originally, these phylogenies contained one cell per subclone; however, we added the remaining cells to the phylogeny as leaves descending from the same node. Phylogenetic correlations (**Methods: Phylogenetic correlations**) were calculated using the one-node depth weighting function. For categorical states (*e.g*., cell type) phylogenetic correlations, weight matrices were first row-normalized before sum normalizing.

To calculate enrichment of heritable genes on each chromosome, the top 2,000 most variably expressed genes (calculated using *Seurat* [Hao et al., 2021]) were segregated by chromosome. Each set of variable genes (on each chromosome) was further divided into genes that were “heritable” (z score *≥* 3) or “non-heritable” (z score < 3). For each chromosome, a Fisher’s Exact test comparing the number of “heritable” and “non-heritable” genes on that chromosome to those on all other chromosomes combined was performed.

### Zebrafish brain development

Normalized RNA matrices and cell annotation tables were downloaded from GEO series GSE105010 and imported into R (v. 4.1.3). Zebrafish [Raj et al., 2018] phylogenies were obtained by parsing json files using code provided by the authors. We used zebrafish 3 (“rep 1”) and 5 (“rep 2”) phylogenies from [Raj et al., 2018]. Phylogenetic correlations (**Methods: Phylogenetic correlations**) were calculated using one-node weighting function, and for categorical states, weight matrices were row-normalized before sum normalizing.

Minor changes were made to the cell annotation provided in the original study. In **Fig 4A** and **Fig 4C**, neuronal cells originally annotated as “S1/S2” (forebrain/midbrain) and “Mix” were both considered as “Mix”. All cell types that were not neurons or neuronal progenitors were considered non-neural.

To impute phylogenetic branch lengths (**Methods: Imputing branch lengths**) for PATH transition inferences (**Methods: Inferring cell state transitions from phylogenetic correlations**), we estimated a cell sampling rate of 10^-4^, which assumes that there were approximately 10^6^ cells per brain [Marhounová et al., 2019].

To classify forebrain neurons as either GABA+, Gluta-matergic (Glut+), or “unassigned”, GABA and Glut marker gene sets were scored across forebrain neuron cells in the rep1 fish (N = 270) using the *Scanpy* [Wolf et al., 2018] `score_genes()` function. Cells with a positive score (greater than 0) for either GABA or Glut marker gene set were classified accordingly (no cells had a positive score for both categories). Cells with scores of 0 in both gene sets were considered “unassigned”.

### Mouse model of pancreatic cancer

Phylogenies, RNA count matrices and phenotype tables were downloaded from GEO series GSE173958 and imported into R (v. 4.1.3). As the available RNA matrices for the murine pancreatic cancer model [Simeonov et al., 2021] were counts, we normalized them using *Seurat* (v. 4.2.0) [Hao et al., 2021]. Also, given that each mouse had been injected with different parental clones whose relationships cannot be established, we could only study the annotated lineages of each clone independently. We analyzed the phylogeny from “Mouse 1 Clone 1” from [Simeonov et al., 2021], which was chosen because it contained the most cells of any clone annotated with an EMT score. All cell annotations were used as published in the original paper. Phylogenetic correlations (**Methods: Phylogenetic correlations**) were computed with the one-node depth weighting function, and for categorical states, weight matrices were row-normalized prior to sum normalizing.

EMT bins were created to discretize the EMT score across the EMT continuum according to the following: cells were partitioned along the continuum using units of 1 (bin #1 includes cells with EMT scores from 0 to 1, bin #2 includes cells from 1-2, *etc*.), merging bins at the extremes (all cells with a score of 7 or lower were assigned to a single bin, as were cells that scored higher than 30) because these bins had low cellular representation. To check for robustness, we repeated the binning procedure using other intervals (0.5,2,3) as shown in **Fig. S5D**.

To impute phylogenetic branch lengths (**Methods: Imputing branch lengths**) for PATH transition inferences (**Methods: Inferring cell state transitions from phylogenetic correlations**), we estimated a cell sampling rate of 10^-6^, which assumes that there were approximately 10^9^ cells per tumor [Del Monte, 2009].

### Human patient glioblastoma

Glioblastoma (GBM) phylogenies and corresponding scR-NAseq data (including gene module scores) were obtained from Chaligne et al. [2021]. Patient sample MGH105 was chosen because tumor location was annotated, and patient samples MGH115 and MGH122 were chosen because each exhibited significant gene module transcriptional heritability in the original paper. The MGH105 phylogeny is a maximum-likelihood (ML) consensus tree, containing 80 cells, 20 cells from each location (MGH105A, MGH105B, MGH105C, and MGH105D). Analyses of patient sample MGH115 used 9 ML phylogeny search replicates for the same 38 cells from the original paper. Analyses of MGH122 used 10 ML phylogeny search replicates and the same 45 cells from the original paper. Phylogenetic correlations were computed by using the inverse node-distance weighting function (**Methods: Phylogenetic correlations**).

PATH inferred transition rates (**Fig. 6F**, **Methods: Inferring cell state transitions from phylogenetic correlations**) were computed using categorical cell states (NPC-/OPC-/AC-/MES-like), with states defined by the corresponding per cell maximum module score, as in Chaligne et al. [2021]. Note that, in the original paper, the NPC-like and MES-like modules combine the NPC1-/NPC2-like and MES1-/MES2-like modules, respectively. PATH inferred transitions 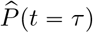 correspond to a time scale proportional to the mean branch length distance separating cells one node apart, *τ*.

Gene set enrichment analysis (GSEA) and Over-Representation Analysis (ORA) were performed using the functions `fgsea()` and `fora()` from the R software package *fgsea* [Korotkevich et al., 2021]. For both analyses, the 3,000 most variably transcribed genes (selected using the `SCTransform()` function from the R software package *Seurat* [Hao et al., 2021] on scRNAseq data) in patient sample MGH115 were ranked by their phylogeny-replicate mean phylogenetic auto-correlation z scores (**Table S4**).

In both analyses, we measured the enrichment of gene sets from the chemical and genetic perturbation (C2:CGP) collection from the molecular signatures database (MSigDB) [Subramanian et al., 2005], as well as the GBM gene modules (NPC1-/NPC2-/OPC-/AC-/MES1-/MES2-like) defined in Neftel et al. [2019], and filtered out sets with fewer than 20 genes. For both analyses (GSEA and ORA), pathway enrichment p-values were adjusted “padj” with the Benjamini-Hochberg procedure (BH), to account for multiple comparisons. Enriched pathways (BH adjusted *p* < 0.05) using GSEA that are presented in **Fig. 6H** were chosen manually (due to putative relevance) from a list of enriched pathways (**Table S5**).

ORA was performed on two gene clusters (“Cluster 1” and “Cluster 2” in **Fig. S6B**), which were determined by hierarchical clustering, using Ward’s method, of the replicate-mean cross-correlations between the top 100 most significantly auto-correlated genes (across the phylogeny-replicates, see **Table S4**) in patient sample MGH115. All 3,000 of the most variable genes were used to define the “universe” or “background” genes to test for over-representation. All enriched gene sets (BH adjusted *p* < 0.05) for Cluster 1, and a manually chosen subset for Cluster 2, are shown in **Fig. S6B**. A complete list of ORA enriched gene sets found in Clusters 1 and 2 from **Fig. S6B** can be found in **Table S6**.

### Gliomasphere phylogenies

Patient-derived human GBM cells (MGG23) [Wakimoto et al., 2011] were grown in Neurobasal Medium (Thermo Fisher Scientific) supplemented with 1/2 x N2 and 1 x B27 (Thermo Fisher Scientific), 1% Penicillin/Streptomycin (Thermo Fisher Scientific), 1.5 x Glutamax (Thermo Fisher Scientific), 20 ng/mL of EGF and 20 ng/mL of FGF2 (Shenandoah Biotechnology). The Molecular Recorder cassette PCT62 [Chan et al., 2019] was introduced into MGG23 cells using piggyBac-mediated transposition (Systems Biosciences). Lineage tracing was initiated by infecting cells with lentivirus expressing Cas9-EGFP, followed by FACS sorting for EGFP-positive cells. Cells were subsequently grown *in vitro* for 4 weeks and lineage traced with the Molecular Recorder approach for two replicates. scRNAseq libraries were generated using the Chromium Next GEM Single Cell GEM, Library & Gel Bead Kit v3.1, Chromium Single Cell Feature Barcode Library Kit, Chromium Next GEM Chip G, and 10x Chromium Controller (10x Genomics) according to manufacturer instructions. Single-cell gene expression libraries were sequenced with paired-end, 28 and 91-base reads on a NextSeq 2000 sequencer (Illumina). The Cas9-edited Molecular Recorder barcodes were PCR amplified from single-cell cDNA libraries as previously described [Chan et al., 2019] and sequenced with paired-end, 28 and 272-base reads on a NextSeq 2000 sequencer (Illumina). Phylogenies were reconstructed using *Cassiopeia* [Jones et al., 2020] using the `VanillaGreedySolver()` with default parameters for each subclone per replicate. ScRNAseq data for each replicate were processed independently using the R package *Seurat* [Hao et al., 2021], by normalizing and scaling RNA count data after subsetting for cells with < 25% mitochondrial DNA and *>* 200 RNA features. GBM gene modules [Neftel et al., 2019] were assigned using the *Seurat* `AddModuleScore()` function. Within each replicate, subclone phylogenies (3 for the first replicate and 6 for the second replicate) were joined at their roots before computing phylogenetic correlations. Phylogenetic correlations were computed for GBM gene modules using the one-node only weighting function, and z scores were computed analytically per replicate. Replicate mean phylogenetic correlation z scores are shown in **Fig. 6G**.

### B-ALL analysis

A blood sample was extracted from a 16yo B-ALL patient after treatment for four weeks with prednisone, daunorubicin, vincristine, and pegaspargase (AALL1131). Rare single persistent blasts were sorted into a 96 well plate based on dim expression of CD45 and CD19 positivity. In addition, CD10, CD20, CD34, and CD38 expression were recorded for each cell. An unsorted remission bone marrow sample was used as a germline control. In addition, a pre-treatment unsorted bulk sample was obtained from the patient at the time of diagnosis. Eighty-six cells with *a priori* tumorigenic phenotype were amplified using primary template-directed amplification (PTA) protocol [Gonzalez-Pena et al., 2021]. Libraries were constructed with the Illumina DNA Prep with Enrichment Kit. All libraries were subjected to whole-exome sequencing at the Chan Zuckerberg Biohub on an Illumina NovaSeq6000. The unenriched libraries were whole-genome sequenced at the New York Genome Center on an Illumina NovaSeq6000 platform. WGS reads were mapped to hg38 using BWA mem and further processed following GATK best practices guidelines [Van der Auwera and O’Connor, 2020]. Somatic single nucleotide variants (SNVs) were detected using an in-house pipeline combining cell genotyping based on *GATK HaplotypeCaller* [Poplin et al., 2017] and somatic detection based on *Mutect2* [Cibulskis et al., 2013]. Cell H3 was removed from the WGS analysis given that it was suspected of being a replicate of H4 because WGS and WES allele frequencies at exonic mutations of H3 did not match. Phylogenetic trees were built with *CellPhy* [Kozlov et al., 2022] using the SNV mutations which were not overlapping with deletions. We detected haplotypic deletions (genomic regions containing only the maternal or only the paternal haplotypes) based on phasing of germline heterozygous SNPs [Delaneau et al., 2019]. Large chromosomal gains were not detected by cytogenetics analyses so we assumed our samples were mainly diploid for the deletion detection analysis. Mutations were mapped to the phylogeny using *treemut* (https://github.com/NickWilliamsSanger/treemut).

The phylogeny was time-scaled using *rtreefit* (https://github.com/NickWilliamsSanger/rtreefit). FACS data were analyzed using the R package *flowCore*. Fluorescence values were compensated and logicle-transformed. Three cells were identified as healthy based on their phenotype, their lower mutation burden and chromosomal deletions, and they were removed from the tree in order to only analyze the tumor population. Fluorescence values were discretized based on frequency using the R package *arules*. Phylogenetic correlations were computed analytically on the discretized fluorescence values using the inverse-node-distance weighting (**Methods: Phylogenetic correlations**). We also classified cells into three states based on the discretized CD19 fluorescence (low: 1-2, medium: 3-4, high: 5-6) and calculated PATH transition rates among those states (**Methods: Inferring cell state transitions from phylogenetic correlations**).

## Supplemental Figures

**Figure S1:**
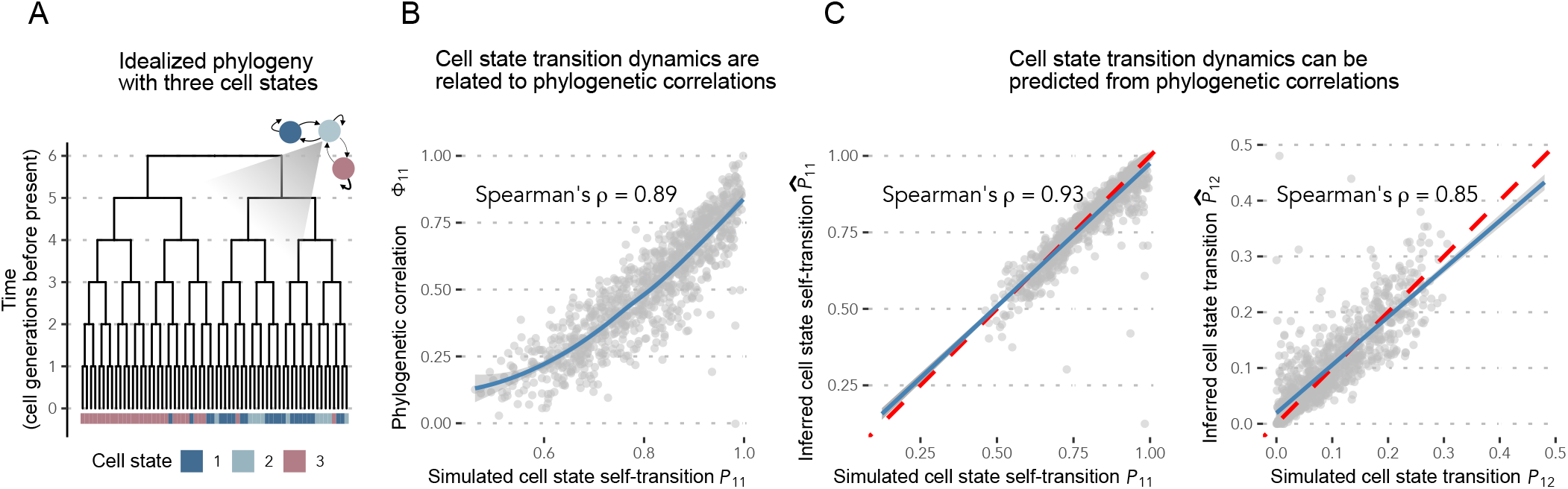
Cell state transition dynamics predict phylogenetic correlations. **A**) Simulated idealized phylogeny containing 2^6^ = 64 cells (**Methods: Simulating phylogenies**) in which cells can transition between three possible cell states. Cell state transitions are represented as a discrete-time Markov chain (**Methods: Markov model of cell state transitions**). **B**) Simulated cell state transition dynamics (**Methods: Simulating phylogenies**) and measured phylogenetic auto-correlations (**Methods: Phylogenetic correlations**) for the first cell state for 1,000 independent simulations on idealized phylogenies, containing 64 cells as in **A**, in which state transition probabilities were randomly generated for each trial. Phylogenetic correlations were computed using a weighting function that included only sister cells (one-node only, as described in **Methods: Phylogenetic correlations and cell state transitions**). LOESS regression line (blue) with 95% confidence interval (light gray) is shown. Spearman’s rank correlation coefficient = 0.89, *p* < 2.2e – 16. **C**) (Left) Simulated versus PATH-inferred (**Methods: Inferring cell state transitions from phylogenetic correlations**), by transforming the phylogenetic auto-correlations measured in **B**, cell state self-transition (*i.e*., stability) probabilities. Spearman’s rank correlation coefficient 0.93, p < 2.2e-16. (Right) Simulated versus PATH-inferred (**Methods: Inferring cell state transitions from phylogenetic correlations**) cell state transition probabilities from state 1 to 2, on idealized phylogenies (**Methods: Simulating phylogenies**). Spearman’s rank correlation coefficient 0.85, p < 2.2e-16. Dashed red lines both have slope 1 and pass through the origin. Linear regression lines (blue) with 95% confidence intervals (light gray) are shown for both plots.

### Box S1: Cell state transition dynamics and phylogenetic correlations

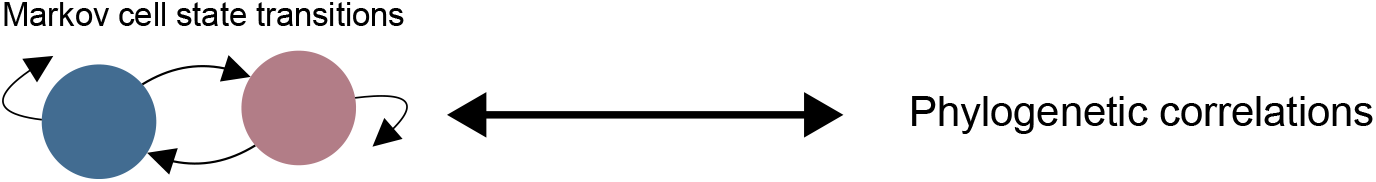 We can connect cell state transition dynamics (*P^t^*) to phylogenetic cell state pair frequencies *F*(*t*), for a given ancestral relationship *t*(e.g., sister cells [*i.e*., *t* = 1] or first-cousins [*i.e*., *t* = 2]) with, (*P*^*t*^)^*T*^*D P* ^*t*^ = *F*(*t*), where *D* = diag(*μ*), is the diagonal matrix of cell state frequencies, and *T* signifies the matrix transpose. This relation, for two cell states, is illustrated below. 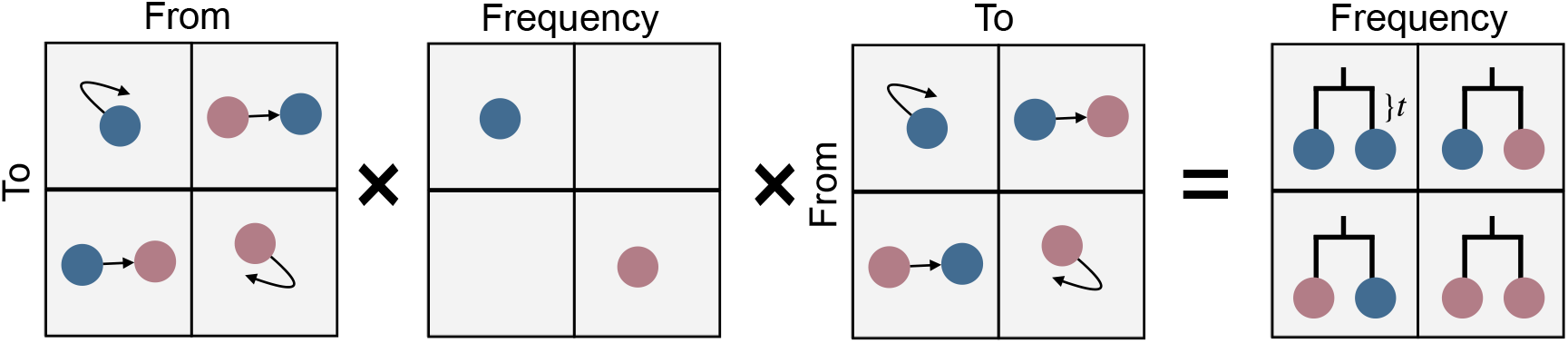 For reversible Markov dynamics, this mathematical relation simplifies to, *DP*_2*t*_ = *F*(*t*). 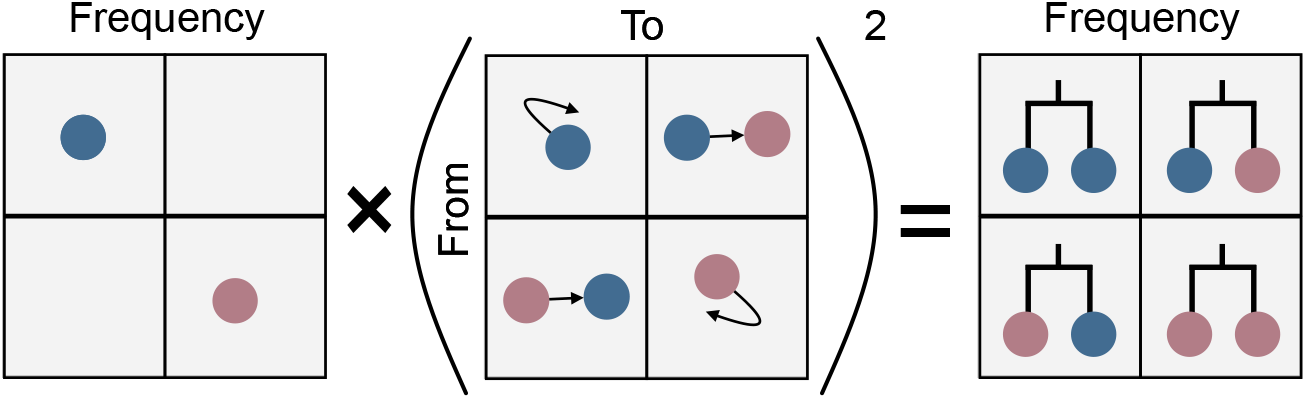 State pair frequencies can be transformed into phylogenetic correlations *Φ*(*t*), by standardizing: *Φ*(*t*) = (*F*(*t*) - *μμ^T^*)/(*σσ^T^*), with *σ*^2^ = *μ* - *μ*^2^. 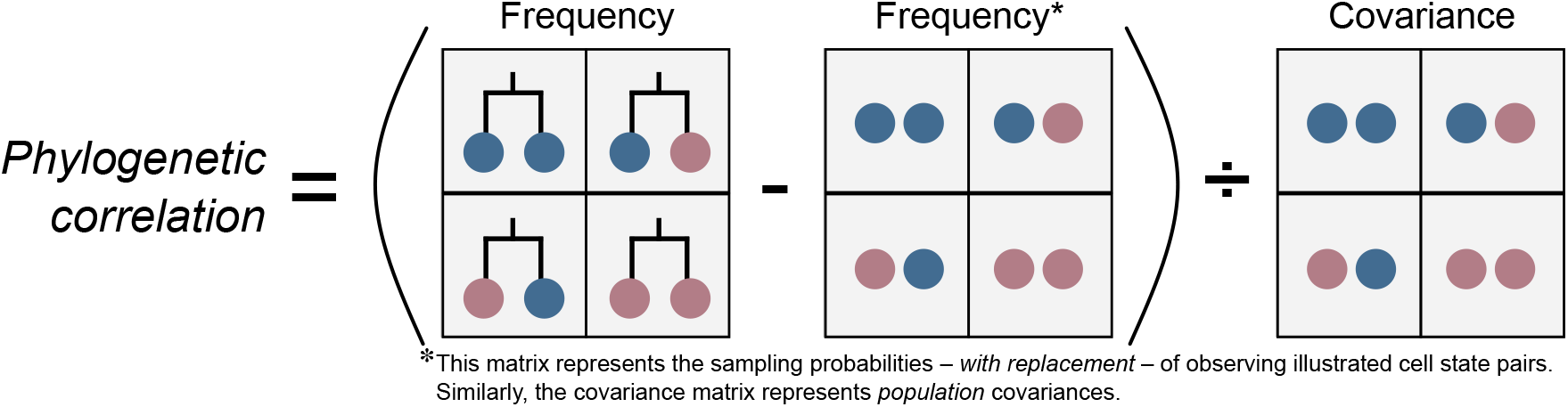 Finally, for reversible dynamics, state transitions can be directly inferred from state pair frequencies, *P*^2t^ = *D*^-1^*F*(*t*).

**Figure S2:**
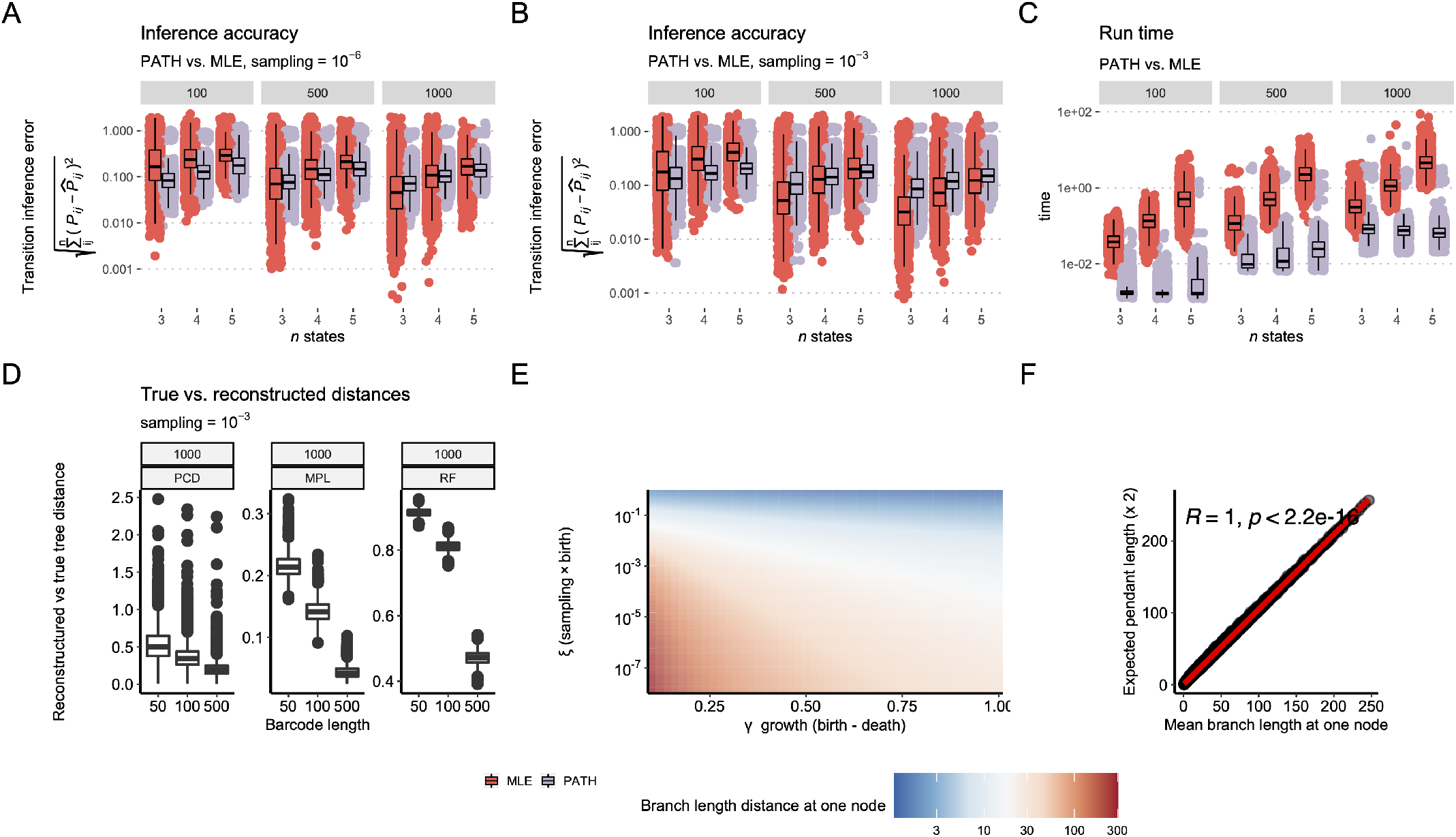
PATH inferences and simulations of somatic evolution. **A**) Transition inference error (Euclidean distance between inferred and true transition probabilities) using PATH or MLE for 3, 4, or 5 cell states in a phylogeny composed of either 100 (left), 500 (middle), or 1,000 (right) cells, representing a sample of 10^-6^ of the total population. Each parameter combination was simulated 1,000 times and inferences are shown for all simulations in which neither PATH nor MLE inference failed. **B**) Same as **A** but with a sampling rate of 10^-3^. **C**) Run times corresponding to simulations depicted in **A**. **D**) Phylogenetic correlation difference (PCD, left), Mean Path Length distance (MPL) [Steel and Penny, 1993] (center), and Robinson-Foulds distance (RF) [Robinson and Foulds, 1981] (right) between simulated true and reconstructed phylogenies (**Methods: Phylogenetic reconstruction**). Phylogenies were simulated 1,000 times for each barcode length (x-axis). **E**) Expected pendant edge lengths for a sampled somatic evolutionary process, as a function of birth, death and sampling rates (**Methods: Imputing branch lengths**). **F**) Correspondence between simulated branch lengths at a node depth of one and expected pendant lengths, while varying sampled somatic evolutionary process parameters.

**Figure S3:**
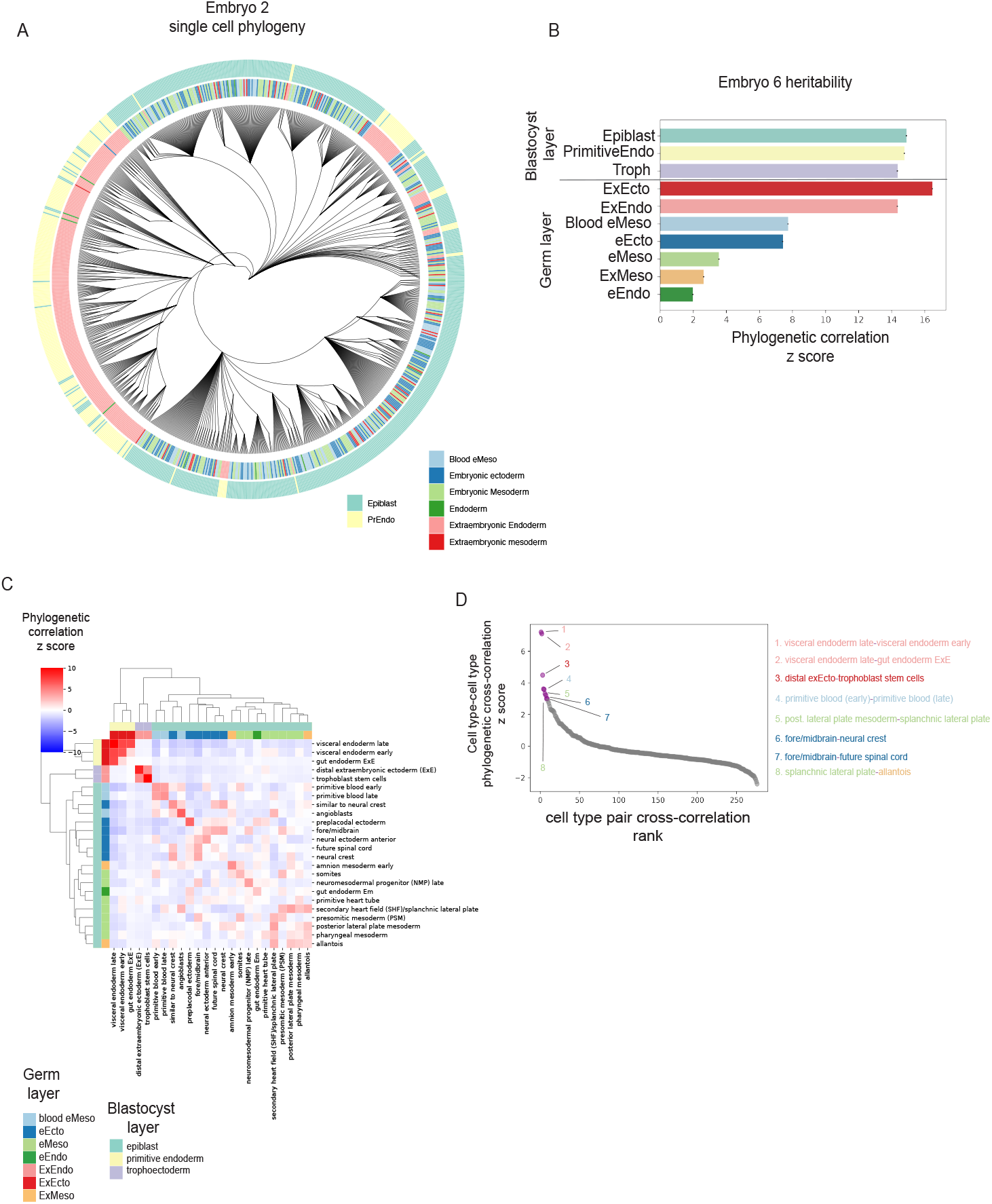
PATH quantifies ancestry and divergence of germ layers and cell types during mouse embryogenesis. **A**) Single-cell phylogeny for mouse embryo 2 from Chan et al. [2022], containing 700 of 1,113 randomly chosen cells for visualization. Each leaf represents a single cell. Leaves are colored by their assignment to a blastocyst or germ layer of origin based on transcription profiles. e prefix, embryonic; ex prefix, extraembryonic. PrEndo, primitive endoderm. **B**) Blastocyst and germ layer phylogenetic auto-correlations for embryo 6 (N = 1,722 cells). **C**) Hierarchical clustering of tissue types in embryo 6 by phylogenetic correlation using Ward’s method. Only tissues with more than 30 cells present in the sample were considered for analysis. Tissues colored by their germ layer and blastocyst layer of origin. ExE, extraembryonic; EM, embryonic. **D**) Ranked pairwise cell type phylogenetic correlations (z scores) for embryo 6. Pairs with z scores > 3 highlighted. Text colored by germ layer as in **B**.

**Figure S4:**
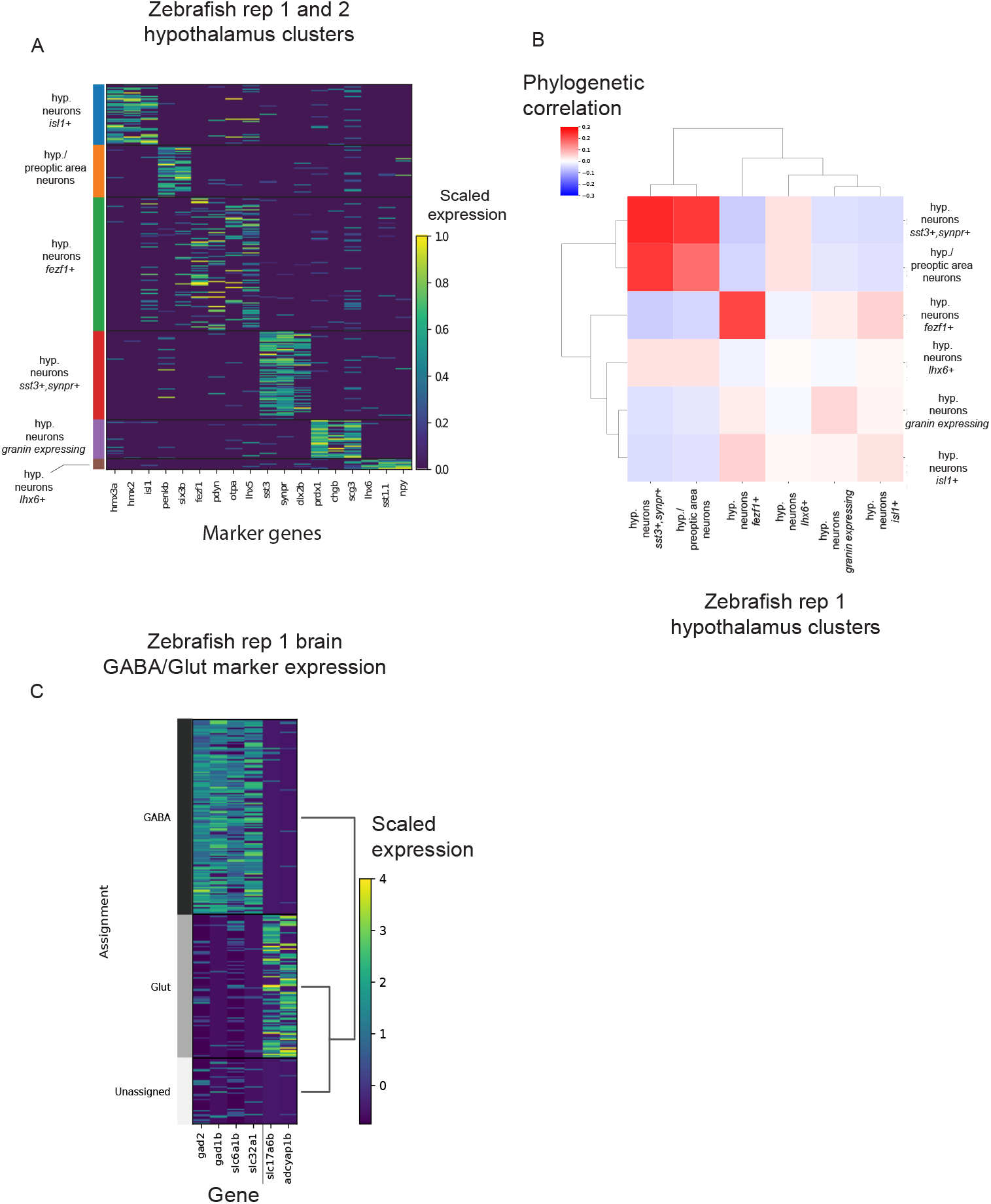
PATH identifies cell fate-determining factors across anatomical, defined tissue and gene expression layers during neurogenesis in zebrafish. **A**) Heat map of scaled expression of representative marker genes across hypothalamus clusters. Marker genes and clusters were defined by Raj et al. [2018]. **B**) Hypothalamus cluster (from **A**) phylogenetic correlations. **C**) Heat map of GABA markers (*gad2, gad1b, slc6a1b, slc32a1*) and Glut (*slc17a6b, adcyap1b*) signaling in forebrain neurons of zebrafish replicate 1 (see **Methods** for assignment of cells into GABA, Glutamatergic (Glut) and Unassigned categories).

**Figure S5:**
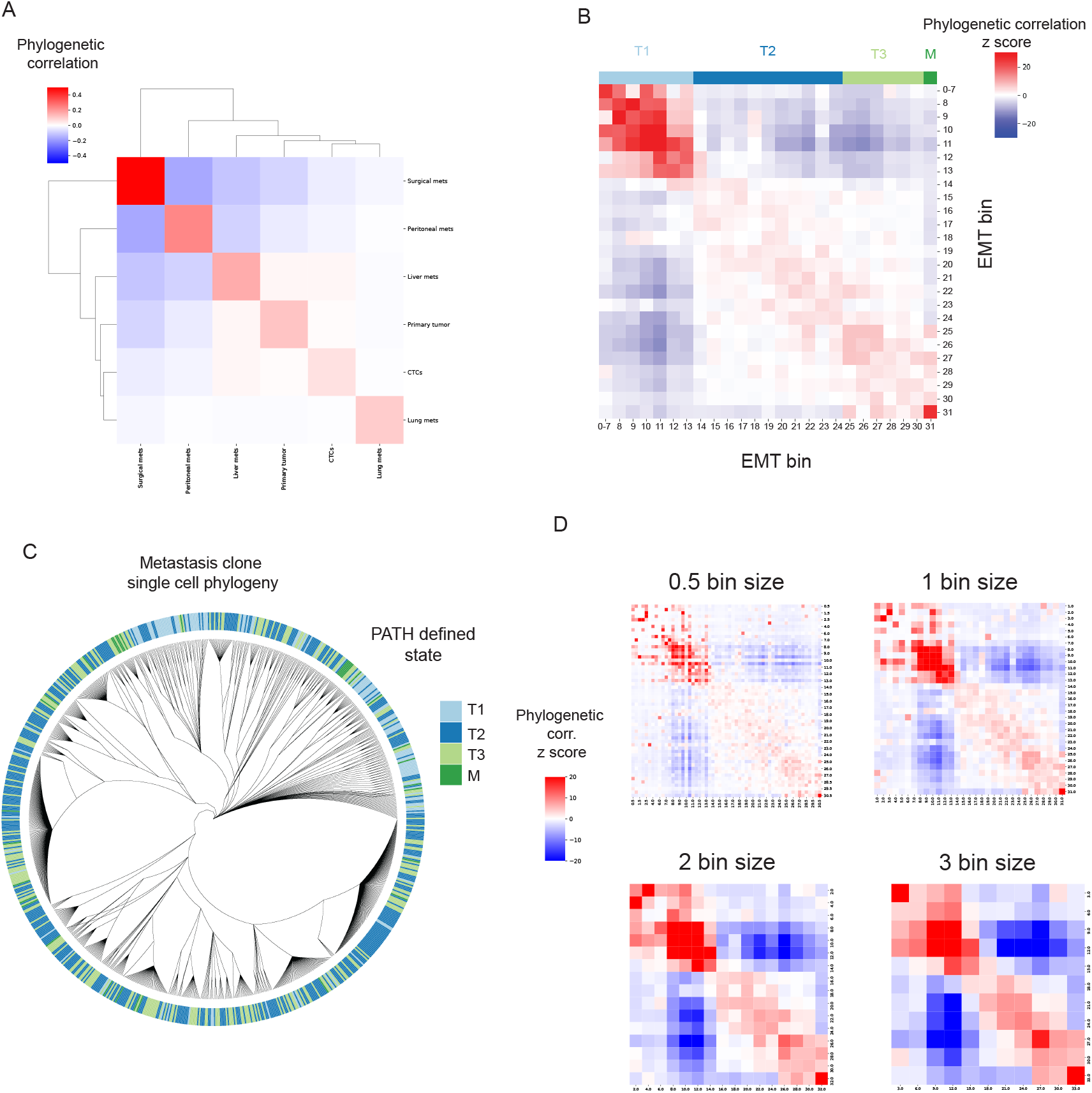
Quantifying the heritability versus plasticity of EMT transcriptional states. **A**) Tumor cell harvest site phylogenetic correlations. **B**) EMT bin phylogenetic correlations (z scores). Colors represent putative states. Full table of EMT bin phylogenetic correlations of can be found in **Table S3**. **C**) Single-cell phylogeny from mouse 1, clone 1 from Simeonov et al. [2021], containing 700 of 7,968 randomly chosen cells for visualization. Each leaf represents a single cell. Cells are colored by PATH-defined states (T1, T2, T3, M). **D**) EMT bin phylogenetic correlation (z score) heat maps using different bin sizes (0.5, 1, 2, 3).

**Figure S6:**
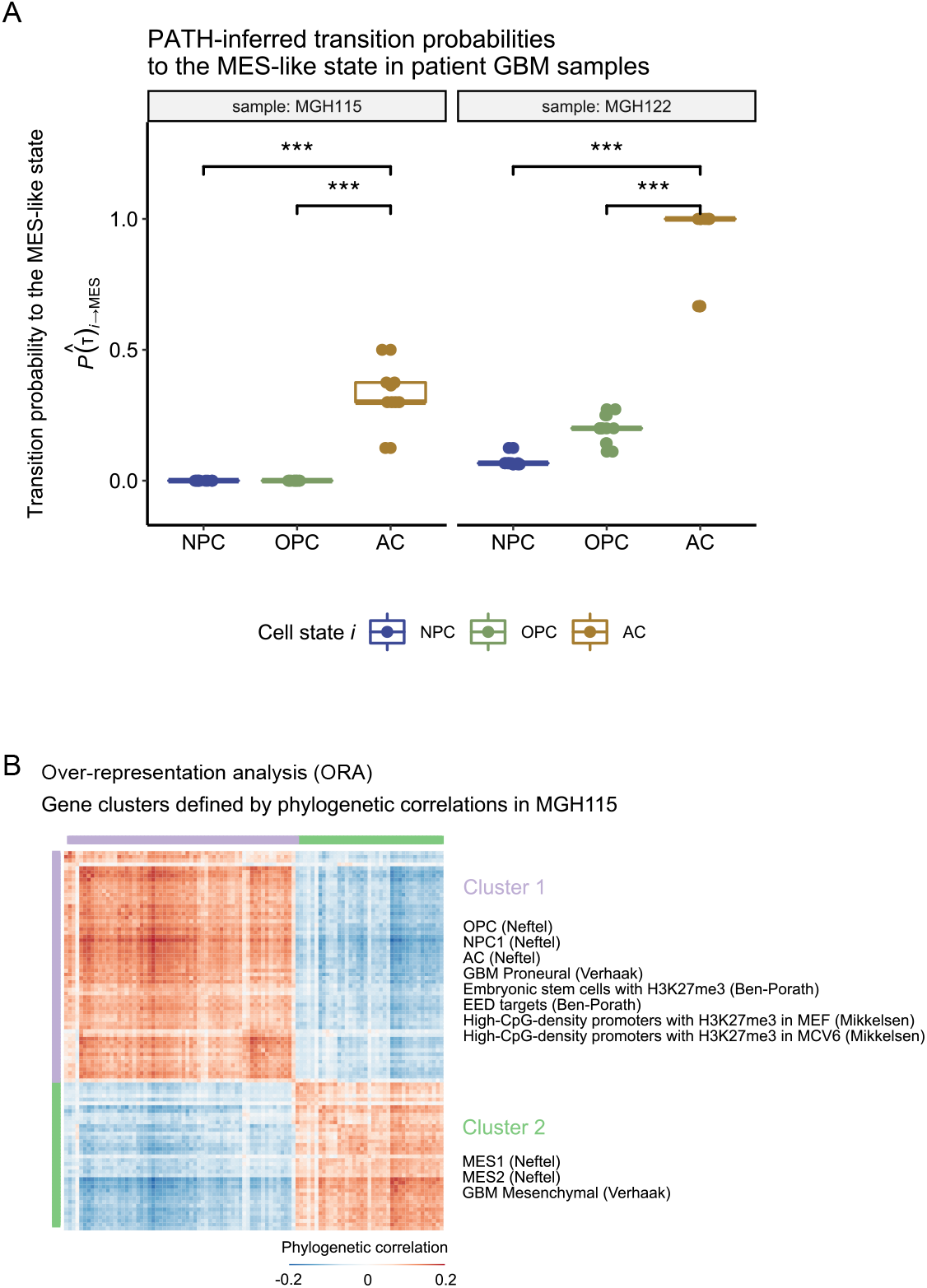
PATH inferred cell state transitions and gene set enrichment in human glioblastoma. **A**) PATH-inferred transition probabilities 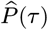 (**Methods: Inferring cell state transitions from phylogenetic correlations**) from neurodevelopmental-like (NPC-/OPC-/AC-like) cell states to the MES-like cell state in human patient-derived GBM samples MGH115 and MGH122 (**Methods: Human patient glioblastoma**). Points correspond to PATH inferences for each sample phylogeny-replicate per sample. Significance determined by two-sided t-test (p < 9.7e-6 and p < 8.2e-9 for NPC-like vs AC-like in MGH115 and MGH122 respectively; p < 9.7e-6 and p < 7.8e-9 for OPC-like vs AC-like in MGH115 and MGH122, respectively). Colors correspond to cell state. **B**) Heat map of the phylogeny-replicate mean phylogenetic correlations (**Methods: Phylogenetic correlations**) for the top 100 most heritable genes (determined by phylogeny-replicate mean gene phylogenetic auto-correlation z scores) in MGH115. Over-representation analysis (ORA) performed on the genes in each of the two clusters, defined by hierarchical clustering using Ward’s method, separately. Phylogenetic correlations were computed using an inverse-node-distance weighting (**Methods: Human patient glioblastoma**). Only select gene sets are depicted for Cluster 2; remaining significantly enriched gene sets are in **Table S6**. GBM gene modules (NPC-/OPC-/AC-/MES-like) were shortened to (NPC/OPC/AC/MES).

**Figure S7:**
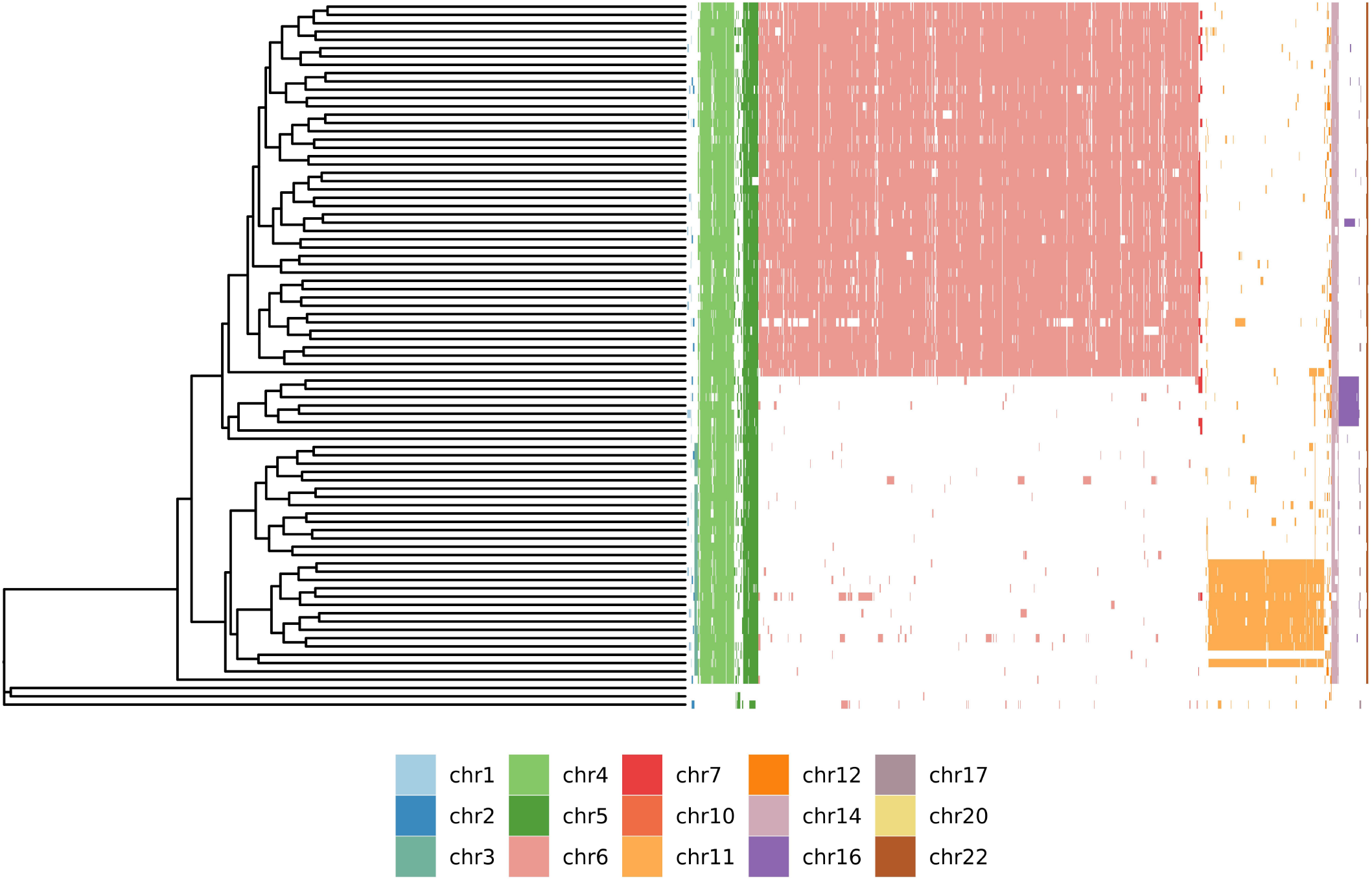
Quantifying cell state heterogeneity in B-ALL using single-cell whole genome sequencing. Genome-wide copy-number deletion annotations projected onto the B-ALL single-cell phylogeny from **Fig. 7A**.

